# Inferring sperm whale (*Physeter macrocephalus*) sex and developmental stage using aerial photogrammetry

**DOI:** 10.1101/2025.10.17.683129

**Authors:** Ana Eguiguren, David Gaspard, Christine M. K. Clarke, Hal Whitehead

## Abstract

Demographic data (i.e. sex and age) are fundamental for analyzing behaviour patterns and evaluating the reproductive potential of a population. However, determining these traits in the wild can be challenging, particularly for marine animals with concealed genitals that spend most of their time underwater. Here, we developed a minimally invasive method to infer the developmental stage and sex stage of sperm whales (*Physeter macrocephalus*) off the Galápagos Islands (N = 51) using uncrewed aerial vehicle (UAV) photogrammetry. We leveraged historic whaling data on sperm whale growth and sexual dimorphism to assign developmental stages to individuals based on their body lengths. We estimated the probability that individuals were female using Bayesian theory based on their morphometry allowed confident classification of the developmental stage and sex for most individuals. Moreover, an examination of the inferred developmental stage and sex individuals that participated in peduncle diving revealed patterns congruent with previous findings that show that this behaviour is predominantly directed at females and performed by subadult individuals. Our method offers an efficient, low-cost means of obtaining demographic information from live sperm whales, contributing to a deeper understanding of the behavioural development and informing population status and viability assessments.

## 1 INTRODUCTION

Demographic data (i.e., sex and age) provide a key dimension for understanding behaviour and evaluating reproductive status in a population. At an individual level, animals of different developmental stages and sexes may adopt distinct social and ecological behaviours that reflect differences in their reproductive strategies and metabolic needs^1–3^. From a population conservation standpoint, knowledge of the demographic structure is essential for assessing life history parameters and their changes over time^4^. In some cases, sex and developmental stages can be easily discerned in the field based on body size, appearance, and behaviour—such as categorizing small, young developmental stages like newborns and determining sex in mature individuals in sexually dimorphic species. However, physically immature individuals can be hard to distinguish from mature ones, and the sexes may be hard to tell apart at this stage. Thus, identifying developmental stages of live animals has often required tracking individuals over time or implementing hormonal analyses^5^, both of which can be invasive and financially and logistically challenging. Likewise, sexing live individuals in the absence of obvious sexual dimorphism can involve genital inspection, which may not be feasible for some species, or relatively costly/invasive molecular analyses of samples collected using variably invasive techniques^6^.

Identifying sex and developmental stage is particularly hard in wild cetaceans, as their bodies are often submerged, which makes visually based assessments impractical. As a result, studies of cetaceans in the wild frequently classify individuals into coarse developmental classes without sex distinctions (calves, juveniles, and adults), or, in species with male-biased sexual size dimorphism, such as sperm whales, into developmental/sex classes that lump together immature males with mature females (e.g. ^7,8^).

The emergence of uncrewed aerial vehicles (UAVs) has allowed researchers to extract precise morphometric measurements of free-ranging cetaceans with minimal impact on their behaviour and wellbeing^9–11^. More recently, UAV-derived morphometric measurements have been used to delineate age classes^12^ and the reproductive status of wild cetaceans^13–15^.

Sperm whales (*Physeter macrocephalus*) off the Galápagos Islands have been the focus of a multi-decade research project spanning 1985 - 2023^16,17^. Because sperm whales in the region are highly mobile^18^ individuals are only feasible to track over a few days and are rarely re-sighted over several decades, making observation-based assessments of their age impractical. Thus, individuals have been classified into four broad developmental/sex classes: calves are considerably small (ca. < 5.5m) individuals found near other larger whales; mature males are considerably large (ca. > 12 m) individuals; immature bachelor males are other individuals found in small (< 4 individuals) groups; and mature females/immature individuals are all other whales found in larger groups^16,17,19^. Although mature males can be reliably identified in the field, as they can be 40% longer and weigh three times as much as mature females^20^, the distinction between bachelor males, immature individuals and mature females is not clear. As the behaviours of mature females and immature males/females are shaped by different social and ecological processes^21^, this grouping masks important facets of their behaviour and population structure.

Informed by age and sex specific morphological data, we developed a method to infer the developmental stage and sex of sperm whales based UAV-derived morphometric measurements, in the absence of known individual demographic data. We first defined size ranges corresponding to finer-scale developmental stages using existing data on sperm whale growth and length-age relationships derived from analyses of thousands of individuals killed during industrial whaling. To distinguish individual sex, we relied on the male’s extreme sexual dimorphism and particularly their disproportionately larger nose, which (when measured from the base of the skull to the tip of the snout) can account for c.a. 40% of their total length, compared to up to 30% of the females’^22^ (Box 1). Although the hypertrophy of male sperm whales’ noses is most notable when they reach physical maturity (> 20 years), it can be detectable in older juveniles (ca. 2 years – 6 m) and intensifies with age^23^. We developed a model-optimizing algorithm to estimate the probability that individuals are females based on their total body length and nose-to-body ratio. To demonstrate the application of our methods, we explored individuals’ involvement in *peduncle dives*—a stereotyped interaction which has thus far been reported only between calves/juveniles and females—in light of our developmental stage/sex class inferences.

## 2 METHODS

### 2.1 Data Collection

We carried out dedicated surveys in the deep waters (> 1000 m) off the Galápagos Islands aboard a 12 m sailboat (*Balaena*) between January and May 2023 (Galápagos National Park research permit No. PC-86- 22). We searched for sperm whales visually during daylight hours and acoustically (using a 100-m towed hydrophone). When we encountered groups of females and juveniles, we followed them for as long as possible at a cautious distance (c.a. 50 m) to collect behavioural, acoustic, and photo-identification data.

If conditions were adequate (wind speed < 10 kt and no rain), we conducted 1 – 2 hour flight sessions, composed of a series of consecutive flights, using a DJI Mini 2 drone (249 g) equipped with propeller guards and landing gear. We conducted sessions when glare from direct sunlight on the water interfered the least with visibility, often early in the morning or later after noon. Over the whales, we flew between 15 - 120 m above the water and pointed the camera down perpendicularly (i.e., nadir). During flights, we recorded continuously at a resolution of 1080 x 1902 or 3840 x 2160 px (4K) resolution, both at 29.79 fps. We alternated between a group-follow protocol–during which we kept visual contact with a group of whales by flying high enough to fit all whales in the frame^24^–and brief moments of close approach (15 - 20 m)–to capture individual whales’ distinctive marks and allow for more accurate size estimates. At the end of most flights, we hovered over the research vessel to collect a calibration image (see 2.2.1 |).

### 2.2 Morphometric measurements

#### 2.2.1 Estimating and correcting measurement error

Errors in aerial photogrammetry arise from several sources, of which the most impactful is altitude measurement error, which impacts the scaling factor used to estimate true object lengths^9–11,25^. Our morphometric estimates were based on altitude measurements from the built-in barometric altimeter, which is less precise (i.e., has more variation) and less accurate (i.e., approaches true values less often) than measurements obtained from laser-based altimeters that are typically used in cetacean photogrammetric research^9,10^. We did not incorporate a laser altimeter on our UAV as it would have exceeded the payload capacity of the airframe and significantly reduced flight time, given the UAV’s small size. We opted for this UAV model because our initial attempts to fly and retrieve a larger UAV (Phantom 4 Pro) equipped with a laser altimeter from the research vessel were largely unsuccessful due to a combination of pilot inexperience (AE) and the inherent challenges of landing a drone on a sailboat at sea. As our main interest was to collect video recordings for analyzing sperm whale behaviour, we opted for the UAV system with reduced of accuracy and precision in altitude measurement in exchange for longer flight times, reliable retrieval, and relatively low replacement cost.

To quantify the uncertainty in morphometric measurements and correct barometric altitudes of our UAV system, we used measurements of our research vessel (12.03 m) collected throughout the field season at various altitudes (27 – 120 m). We quantified percent measurement error using a modified version of the equation in Bierlich et al. (2021)^10^:

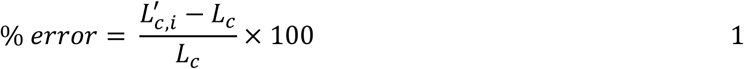

Where 𝐿_𝑐_ is the known length of the calibration object in meters, and 𝐿^′^ is the estimated length in meters of the calibration object in each image 𝑖. We used MorphoMetriX V2^26^ to measure the length in pixels (𝐿_𝑐𝑝_) of the research vessel in still images taken from video recordings, and converted length measurements in pixels (𝐿_𝑐𝑝_) to length (𝐿_𝑐_) in meters by applying equation (2), modified from Burnett et al. (2019)^9^:

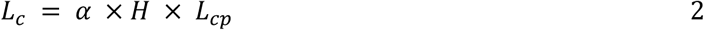

where *H* is the drone altitude above sea level, and α is a scaling factor corresponding to the DJI Mini 2 drone camera. While 𝛼, can be computed based on known camera parameters (i.e., focal length and pixel dimensions), these values were unavailable for our drone model from the manufacturer. We therefore empirically estimated α by obtaining 𝐿_𝑝_ measurements of a known object of known length *L* at a known distance (*H*) in the lab and then using equation (3).

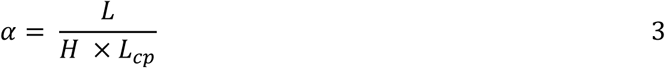

To estimate the bias in the drone’s barometric altitude, we first computed the true altitude 𝐻_𝑇_ given the 𝐿_𝑐𝑝_ for each still image of the research vessel and its known length 𝐿_𝑐_.

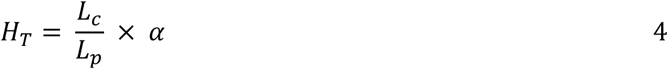

We then applied a linear regression to estimate a corrected altitude (𝐻_𝑐_) given the barometer altitude:

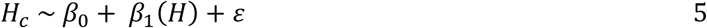

To account for the possibility that barometric altitude biases would vary on different days as a result of changes in weather conditions, we also fit random effects models with date as a random intercept and slope. Although we found evidence for variation in the intercept and slope across different dates, this had a negligible effect on measurement error. This likely results from our study area having very little variation in barometric pressure within and between days. The results of random effects models including date are shown in **Supplementary Material 1**.

#### 2.2.2 Measuring whales

Drone footage was quality-rated on a scale of 0 – 8, with 0 being high quality and 8 being low quality, based on the level of glare, sea-surface disruption, focus, and exposure (described in **Supplementary Material 2**). Only recordings with a quality rating ≤ 4 were included in the analysis. Within high-quality videos, we extracted still images using the behavioural analysis software BORIS^27^. We selected frames where whales were lying mostly flat at the water surface, located near the center of the frame, where the possible effects of lens distortion are negligible^9^, and when the drone camera was positioned at nadir relative to the water surface, as confirmed by the flight log data. We attempted to capture a broad size range of individuals, so we note that measured whales are not a random sample of the population.

For each whale, we measured the total length (*TL*) and two alternative nose length measures—snout-to- flipper length (*SnF*) and snout-to-dorsal-fin length (*SnD*)—in pixels (Figure 1). *TL* was measured piecewise from the snout to the fluke notch, *SnF* was measured from the snout to the transversal intersection of the anterior base of the flippers with the spine (when at least one flipper was visible), and *SnD* was measured from the snout to the caudal base of the dorsal fin. To estimate nose proportions, we calculated the nose-to-body ratio (*NR*) by dividing *SnF* or *SnD* by *TL* (in pixels), resulting in two metrics: *NR_flipper_* and *NR_dorsal_,* respectively. *TL* was converted from pixels to meters using Equation 2, incorporating the corrected drone altitude (𝐻_𝑐_) calculated using Equation 5.

**Figure 1.**
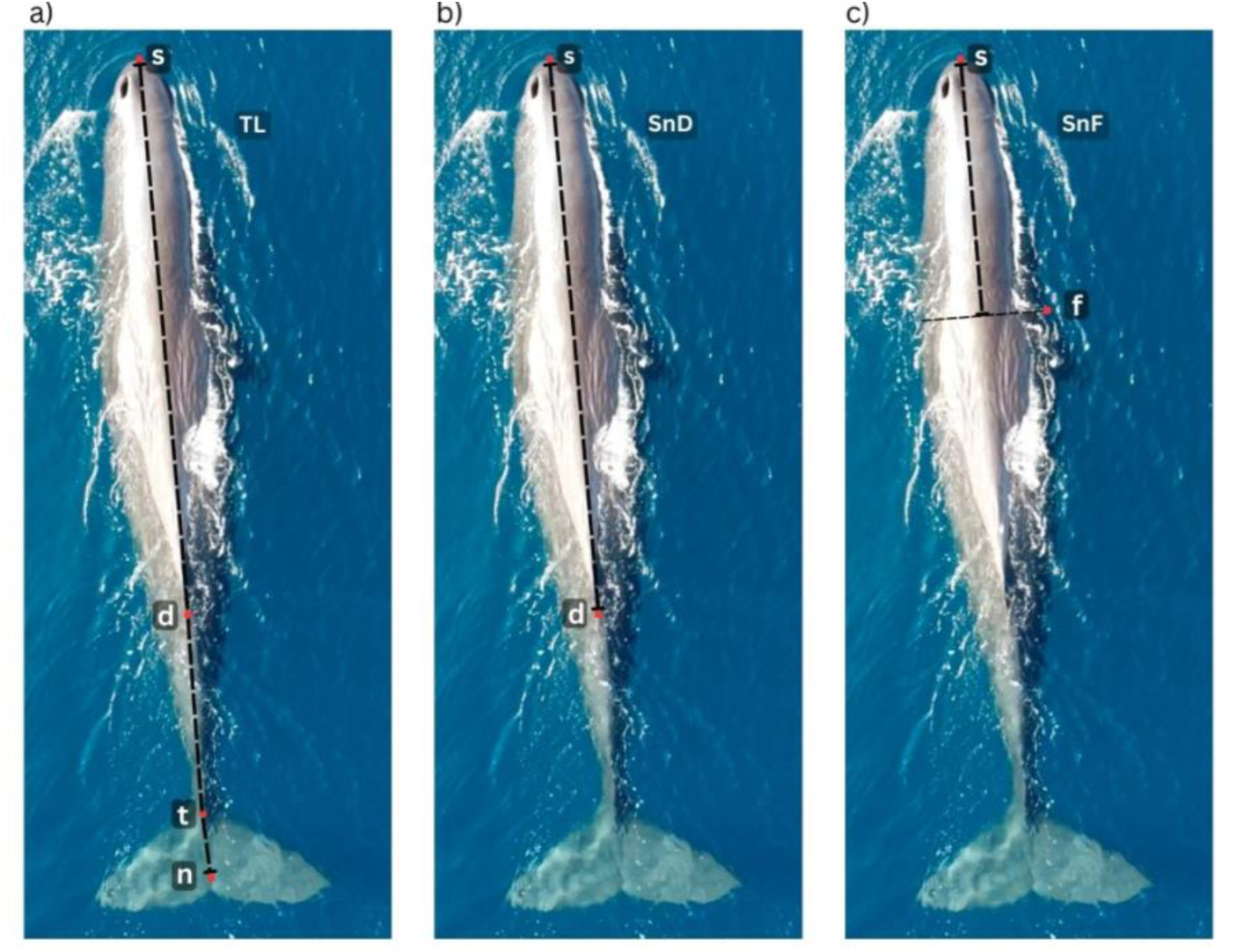
Aerial images of a sperm whale showing landmarks used to measure sperm whale morphometry (s = snout; f = flipper insertion point; d = dorsal fin; t = tail-stalk; n = fluke notch). Total length (*TL*) measures the piece-wise distance from s to d, to t, to n (a). Snout-to-dorsal fin length (*SnD*) measures the distance from s to d (b). Snout-to-flipper length (*SnF*) measures the distance from s to the midpoint of the spine that intersects perpendicularly with f (c).

To capture inter-image variability, we aimed to measure each whale at least three times per recording. However, obtaining *SnD* and *SnF* measurements was sometimes hindered by whale positions and visibility. As sperm whales often tuck their flippers against their body, the insertion point of the flipper could not always be observed from the drone’s perspective, which impeded measuring *SnF*. *SnD* measurements were limited by light and water conditions, or when the dorsal fin gradually tapered into the body without a clear boundary. To compare the variability across images, we obtained average coefficients of variance (% CV), calculated by dividing the standard deviation (SD) by the mean x 100 for measures taken from the same individual.

### 2.3 Photoidentification

We identified measured whales from their aerial photographs based on observable markings—including visible fluke marks, scars, indentations, rake marks, white patches, and sloughed skin patterns^28^. We rated images used for measurement on a scale of 1 – 5 (1 = poor, 5 = good) based on focus, contrast, and saturation (modified from ^29^). Initial identifications were made using images rated ≥ 3. In cases where multiple still images of the same individual were taken from a video recording, we also assigned identifications to lower-quality images if contextual evidence supported the match to a higher-quality image (for example, if the same whale could be tracked throughout a recording).

### 2.4 Inferring sex and developmental stage

#### 2.4.1 Sex

The relationship between sperm whale length (*L*) and nose-to-body ratio (*NR*) depicted by Nishiwaki et al.^23^ can be modelled by separate logistic curves for males and females. For females, the nose-to-body ratio (𝑁𝑅_𝐹_) can be modelled as follows:

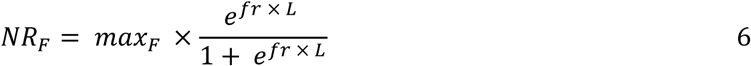

Where 𝑚𝑎𝑥_𝐹_ is the maximum (asymptote) *NR* of female whales, and 𝑓𝑟 is the initial rate of change in *NR* with increasing length. For males, the relationship between body length and *NR* for young (i.e., small individuals) is expected to follow the same trend as that of females, diverging after a length threshold (𝑐ℎ𝑚) such that:

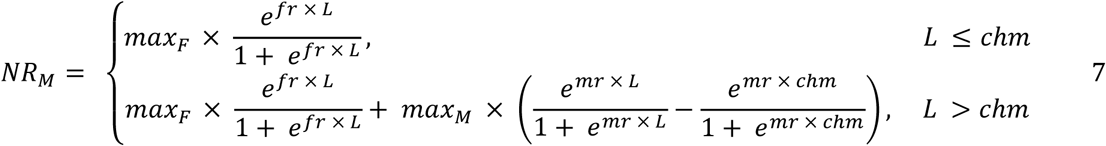

Where 𝑚𝑎𝑥_𝑀_ is the maximum *NR* for male sperm whales and 𝑚𝑟 is the initial rate of change in *NR* with length when length is greater than the threshold (*chm*), which we set at 6 m based on ^23^.

We found the parameter values for *max_F_, fr, max_M_,* and *mr* that minimized the total sum-of-squares given our data, using the *optim* function with the Broyden-Fletcher-Goldfarb-Shanno (BFGS) optimizing algorithm in base R^30^. We initialized the optimization algorithm using parameter estimates based on ^23^’s figure showing the relationship between total body length and *NR* estimates, in which nose length was measured from the tip of the snout to the eyeball. We constrained the optimizing to *mr* values > 0 (i.e., always positive) to ensure that modelled *NR* values based on male growth models would be higher than those of the female growth model.

The posterior probability that each individual was female was estimated based on Bayes’ theorem under the prior assumption that individuals would be equally likely of either sex. Estimates of individuals’ likelihood of being female (𝐿_𝑓𝑖_) were computed based on how close each point fell to the ‘female curve’ following equation (8).

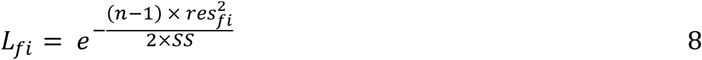

Where *n* is the total number of individuals, *res_fi_* is the residual between an individual’s observed *NR* and that predicted by the female curve, and *SS* is the sum of the residuals (both for the female and male curves). The likelihood of being male (𝐿_𝑚𝑖_) was calculated similarly. We then computed the posterior probability of an individual being a female (𝑃(𝑓_𝑖_)) using equation (9).

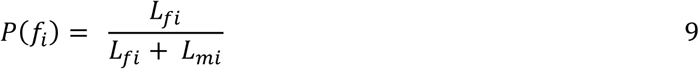

We report our results in terms of *P(f)* but note that the probability of an individual being male under this framework is the direct opposite (*P(m) = 1 – P(f))*.

To propagate the uncertainty associated with inter-image variation in estimates of individual probability of being female, we applied a stratified bootstrap simulation by individual ID^25,31^. In each of 1000 iterations, we randomly sampled a set of measurements from one still image for each individual whale. Sampled data was used to obtain optimized parameter values and individual probabilities of being female. Resulting estimates were then used to calculate mean values and 95^th^ percentile confidence intervals (95% *CI*). For this analysis, we included only individuals with at least three measurements of *TL, NR_flipper_*, and *NR_dorsal_*.

We carried out robustness checks evaluate the effect of our modelling assumptions on individual’s *P(f)* estimates to our modelling decisions. Specifically, we systematically varied *chm* values within a reasonable range (5 – 7 m) to compare resulting posterior *P(f)* estimates. We also computed posterior *P(f)* estimates using a prior *P(f)* set to 0.79, which corresponds to the proportion of females in breeding groups off the Galápagos Islands genetically estimated in 1991^32^.

#### 2.4.1 Developmental stages

We associated individual length (*TL*) to developmental stages defined in **Table 1**, which we delineated based on whaling-based research relating length measurements to analyses of gonadal development, stomach contents, and teeth-layer based age estimates^20,33,34^.

**Table 1.** Developmental stages for male and female sperm whales based analyses of whaling data. ^20,33,34^.

| Age class | Sex | Length (m) | Age range | Life stage traits |
| --- | --- | --- | --- | --- |
| Neonates | Both | < 4.1 | Few days - months | Unhealed umbilical regions, likely recently born<sup>34</sup>. |
| Calves | Both | 4.1 – 5.5 | < 1 year | Young individuals almost exclusively dependent on milk<sup>34</sup>. |
| Juveniles | Both | 5.5 – 7.6 | 1 – 2 years | Primarily depend on milk for sustenance, although solid foods have been found in their gut contents<sup>34</sup>. |
| Sub-Adult | Male | 7.6 – 10.0 | 2 – 7 years | While milk may still be present in the stomach, solid foods are more frequently found. Still, the majority of individuals have not attained sexual maturity<sup>34</sup>. |
| Sub-Adult | Female | 7.6 – 8.5 | 2 – 7 years | While milk may still present in the stomach, solid foods are more frequently found. Still, the majority of individuals have not attained sexual maturity<sup>34</sup>. |
| Adult | Male | 10.0 – 13.7 | 7 – 20 years | Sexual maturity in males (sperm production) starts at 10 m long, between 7 – 11 years of age, which corresponds with them leaving their natal unit<sup>33–35</sup>. |
| Adult | Female | 8.5 – 10.m | 7 – 20 years | Females achieve gonad development between 8.2 – 9.2 m and are able to conceive shortly after<sup>20</sup>. |
| Mature | Male | > 13.7 | > 20 | Almost all males of this size range are physiologically fertile, as defined by the concentration of sperm in their seminal fluid and will be either solitary or form bachelor schools. Although physiologically fertile, males will likely only start mating when they've reached >15.7 m (35 years)<sup>20,34</sup>. |
| Mature | Female | 10 – 12 m | > 20 years | Females attain full size at this age. |

#### 2.4.3 Behavioural context

We then inspected whether individual whales performed or received *peduncle dives* across age classes and differing probabilities of being female in videos where at least 2 whales were observed. *Peduncle dives* are short (a few seconds) and shallow dives usually performed by a calf or juvenile next to the base of the peduncle (fluke stalk) and beneath a larger whale, during which the calf/juvenile often presses its snout onto the larger whale’s genital region^36^. They can be detected on drone-based recordings when calves arch their backs and dive under a larger whale’s body repeatedly. For each measured whale, we recorded whether it had been observed performing or receiving a *peduncle dive* in any of the video recordings from which still images for measurements were extracted.

Peduncle dives have been assumed to indicate suckling or to stimulate the release of milk^36^. Underwater footage suggests that they may not be associated with direct suckling, although the role of this behaviour in promoting milk letdown remains unclear^37^. Alternatively, peduncle dives may represent a form of affiliative behaviour between young whales and mothers/allomothers^37^. Still, although peduncle dives may not necessarily involve suckling, all reports of peduncle dives in which the sex of the receiving whales is known involve females^36–38^.

#### 2.4.4 Exploring performance on an external dataset

To examine the performance our method on whales observed in other populations, we also included morphometric measurements of two individuals, one observed in the North Atlantic (The Gully Canyon off the Scotian Shelf, Canada) and the other in the Arctic Ocean (Baffin Bay, Canada). The North Atlantic individual was recorded using a DJI Phantom 4Pro V2 + and the Arctic individual was recorded with a DJI Inspire 2 equipped with a DJI Zenmuse X5S micro four-thirds camera with Olympus M.Zuiko lens. UAV altitude measurements were obtained through laser altimetry using a LidarBoX system^39^. We computed the posterior probabilities of these individuals being female based on their *TL* an *NR_flipper_* measurements and the optimized parameter values and individual likelihoods of being male or female from our original Galápagos dataset. Because these individuals were located in high latitudes^20,40^ we expected them to have near-zero probabilities of being female.

## 3. RESULTS

### 3.1 Error estimation and correction

We obtained 343 measurements of *Balaena* for calibration across 18 days in the field between February 1^st^ and April 29, 2023, at varying altitudes. Length estimates based on barometric altitudes underestimated the boat length by 0.55 m on average (*SD* = 0.37 m), corresponding to a -4.55% measurement error (*SD* = 3.15%). This bias was associated with an average 2.35 m underestimation in barometric altitude (*SD* = 1.94 m). Using the model corrected altitude (𝛽_0_ = 1.40 (𝑆𝐸 = 0.21); 𝛽_1_ = 1.017 (𝑆𝐸 = 0.003)) reduced average length estimate error to 0.12 % (*SD = 3.15%,* CV = 3.15%).

### 3.2 Whale measurements and photo-identification

We were able to extract *NR_dorsal_* metrics more frequently than *NR_flipper_* (491 and 297, respectively). Only images captured at altitudes of 70 m or less were high enough quality (Q3 – 5) for initial identification (Figure 2), resulting in 504 still images assigned to 90 individuals for which *TL* could be measured at least once, and a subset of 168 still images assigned to 51 individuals for which *TL, NR_dorsal_,* and *NR_flipper_* could be measured at least three times.

**Figure 2.**
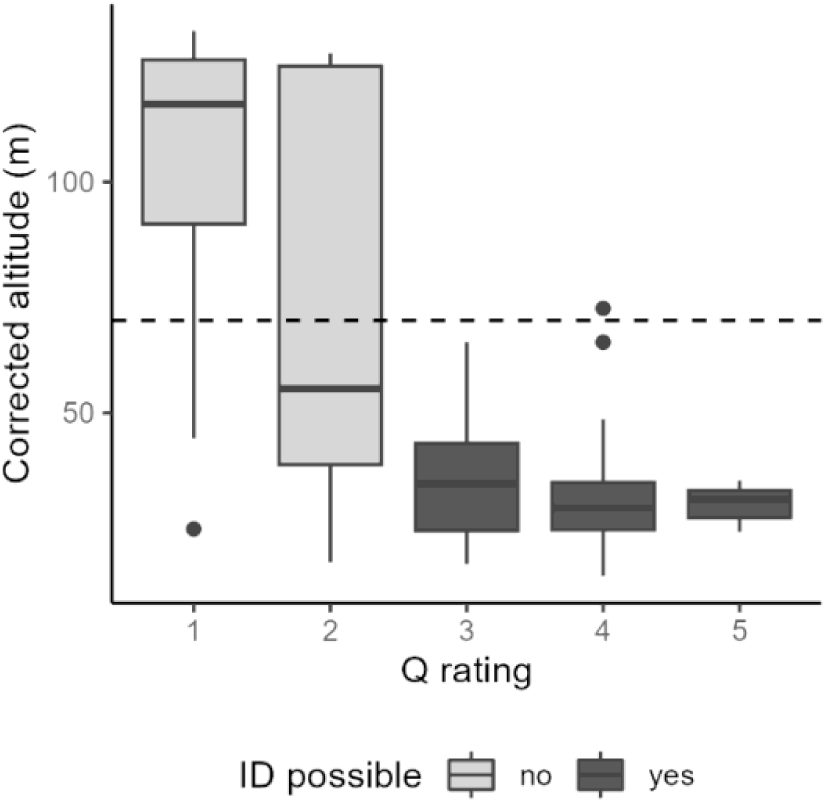
Corrected drone altitude (m) distribution across photo quality ratings (Q1 -5) of still images. The 70 m threshold is shown for reference.

### 3.3 Developmental stage and sex inference

#### 3.3.1 Uncertainty in individual measurements and developmental stage assignments

Observed *TL* measurements of the same individual had an average 2% CV (*SD* = 1.39%). The 95% CI width in bootstrapped estimates of sperm whale *TL* had a median of 0.35 m (mean = 0.42, *SD =* 0.32). This represented a median of 3.29% of the mean *TL* (mean = 4.18%, *SD =* 3.34%). Resulting *TL* estimates ranged from 4.1 -16.1 m, with 80% of individuals measuring between 7.4 – 12.6 m (**Figure 3**). These estimates resulted in no individuals categorized as neonates, three as calves, three as juveniles, one as a subadult, and four as mature males. The remainder (n = 40) fell within age classes with overlapping ranges between males and females (i.e., AF, AM, and MF).

**Figure 3.**
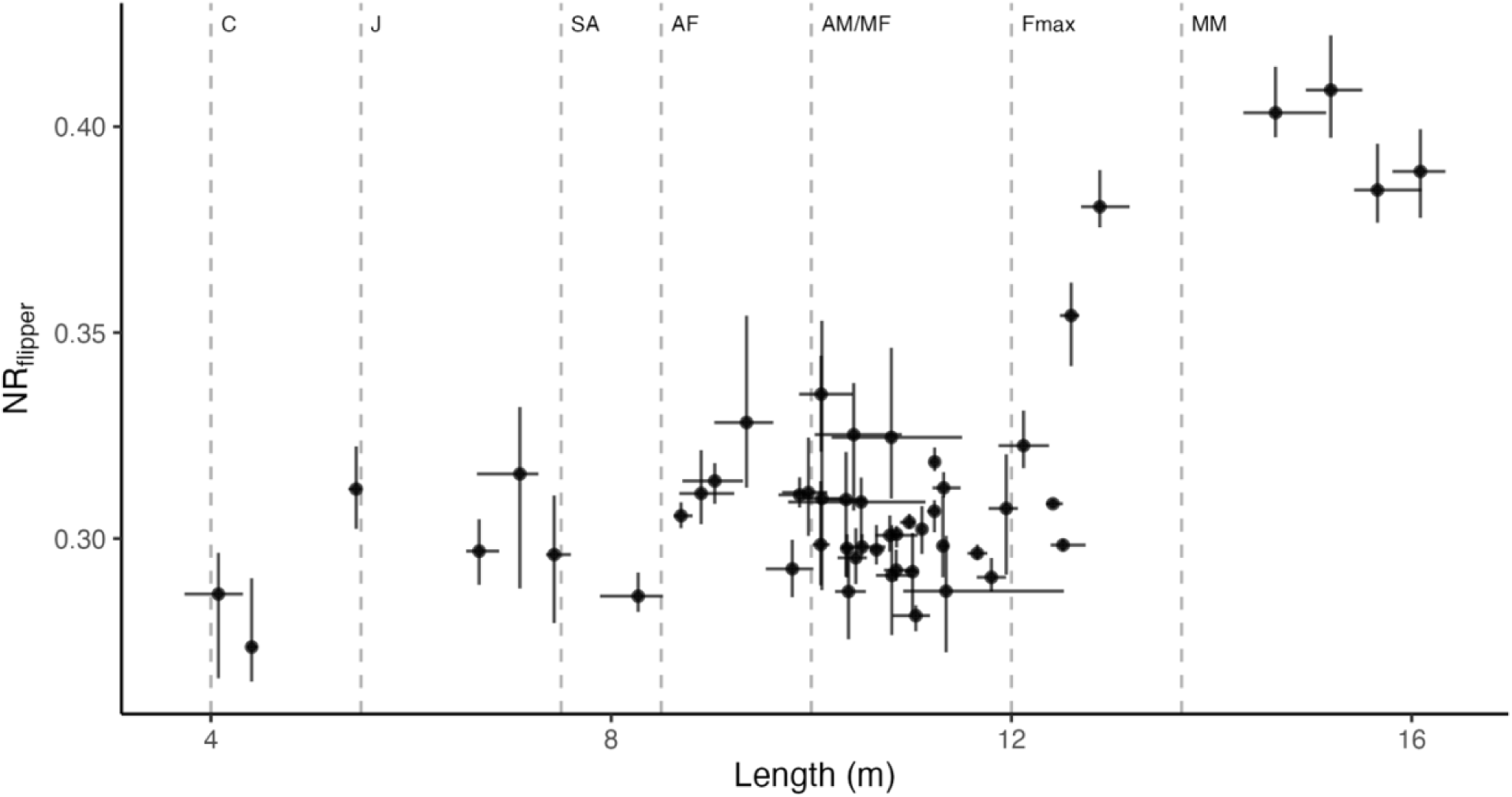
Total body length (m) and nose-to-body ratio (*NR_flipper_*) estimates of individual sperm whales. Point locations show the bootstrapped mean for each individual (N simulations = 1000), horizontal error bars show the corresponding 95% CI length range, and vertical error bars show the bootstrapped 95% CI *NR_flipper_* range. The dashed vertical lines indicate the minimum body lengths associated with sperm whale sex and age defined in **Table 1** as follows: calf (C), juvenile (J), sub-adult (SA), adult female (AF adult male and mature female (AM/MF), maximum female length (Fmax), and mature male (MM).

*NR_flipper_* measures ranged from 0.27 – 0.41 (mean = 0.31, SD = 0.03). On average, *NR_flipper_* measures from the same whale had an observed 2.95% CV (SD =2.12%).

#### 3.3.2 Parameter optimization

We found the divergence between mature males and the rest of the measured whales was much less pronounced for *NR_dorsal_* measurements than *NR_flipper_* measurements, contributing to greater uncertainty in *P(f)* based on this metric. We describe results for models fit using *NR_flipper_* below and those for models fit using *NR_dorsal_* in **Supplementary Material 3**. Optimal *fr* values were variable across bootstrap iterations, resulting in a high degree of uncertainty in modelling the *NR_flipper_*of smaller (< 6 m) whales (**Figure 4** & **Figure 5**). Still, the divergence in *NR_flipper_* between males and females after *chm* was consistently pronounced, partly because large males (> 13. 7 m) had disproportionately higher *NR_flipper_* than the rest of individuals (**Figure 6**). Estimates of asymptote parameters (*max_f_* and *max_m_*) were less variable than growth parameters (*fr* and *mr*) (**Figure 4**). Additionally, for adult males *NR_flipper_* seems to increase linearly with length (**Figures 5**), and thus the logistic model may be an unnecessary elaboration. This is also supported by *max_m_* often estimated to be impossibly greater than 1.0; **Figure 4**).

**Figure 4.**
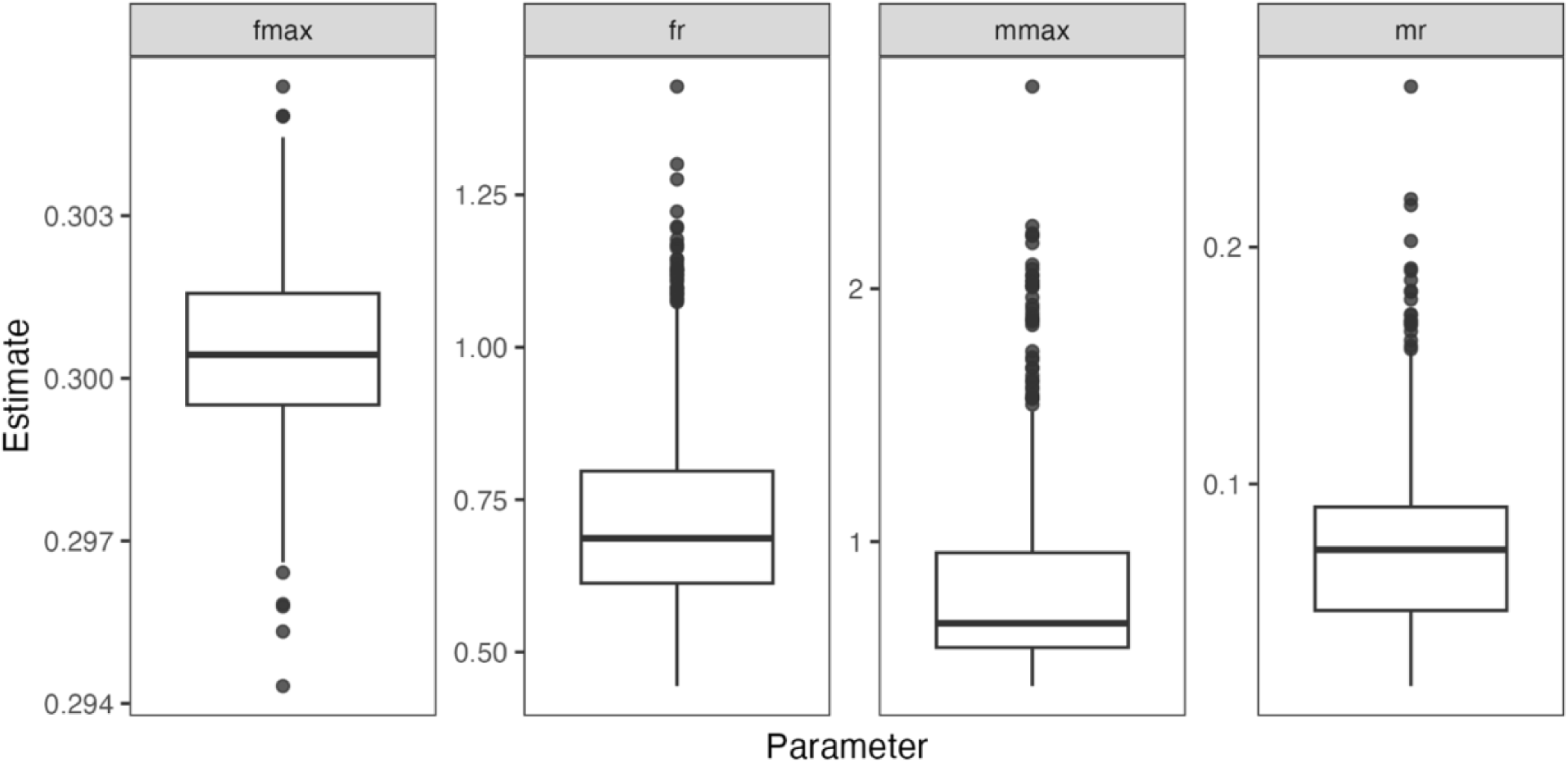
Distribution of bootstrapped parameter estimates modelling the growth rate of females and small males (≤ 6 m) (*fr*), the female asymptote of *NR* (*max_f_*), the growth rate of larger males (> 6 m) (*mr*), and the male asymptote of *NR* (*max_m_*).

**Figure 5.**
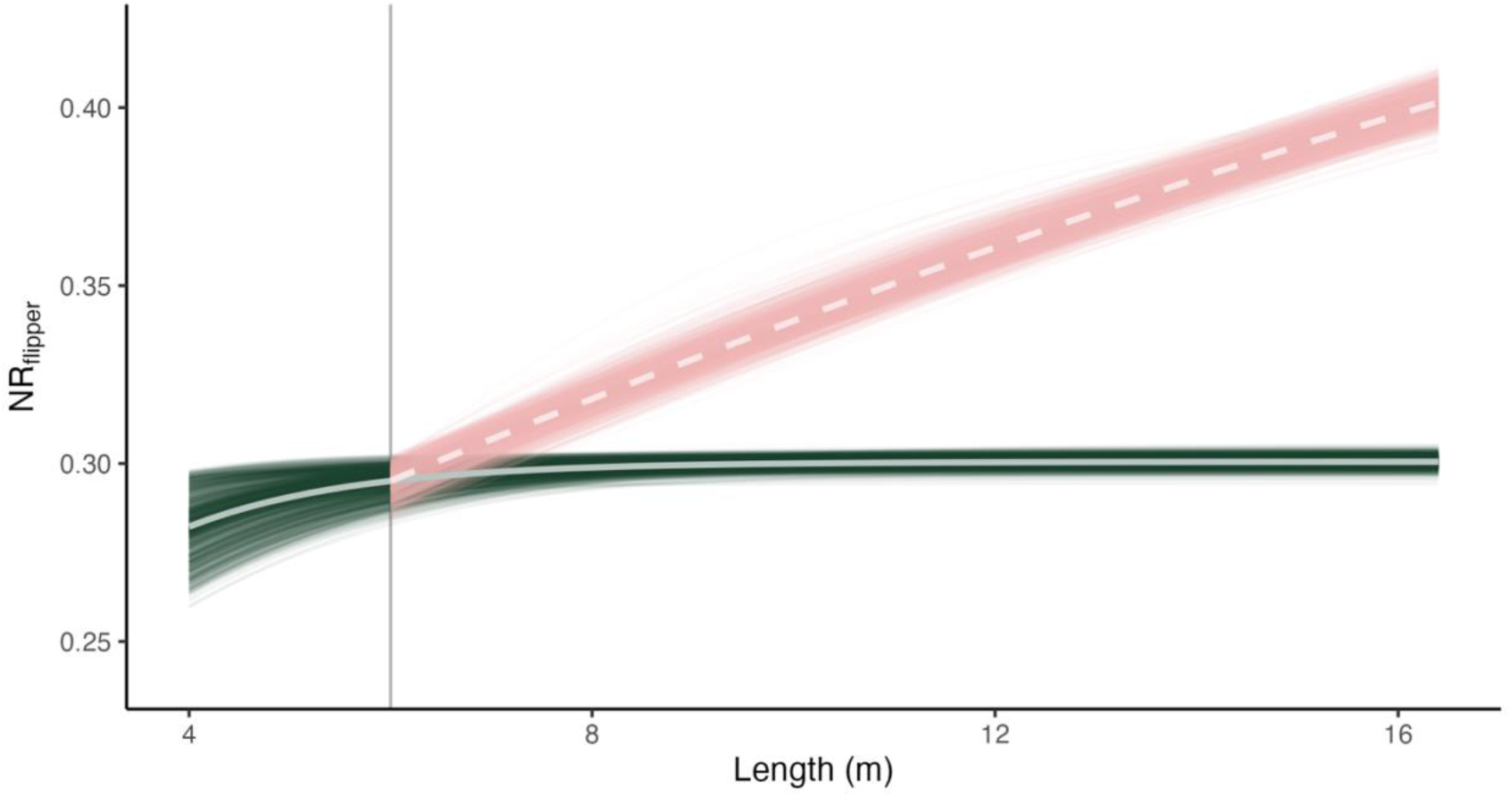
Bootstrapped logistic curves of the total length (m) and the nose-to-body ratio of sperm whales based on measures of the snout to the anterior base of the dorsal fin (*NR_flipper_*_)_. Theoretical male curves are shown in pink and theoretical female curves are shown in green. The average *NR_flipper_* values across iterations are shown by light violet dashed pink and green solid lines for males and females, respectively. The vertical line indicates the point of divergence between males and females (*chm =* 6 m).

**Figure 6.**
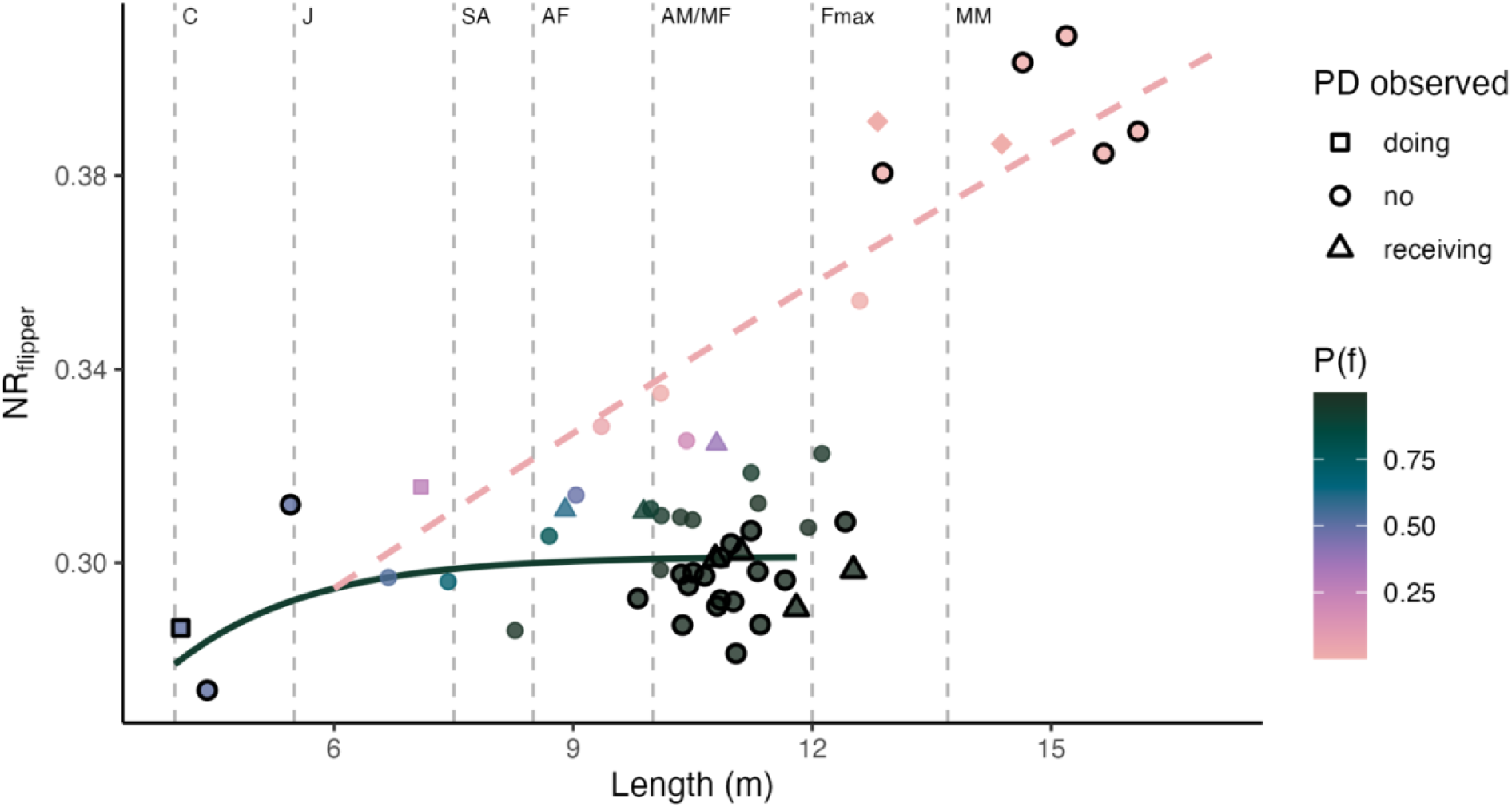
Bootstrapped mean Length (m) and nose-to-body ratio (*NR*) for individual sperm whales based on snout –– flipper distance (*NR_flipper_*). The solid green line and dashed pink line show the modeled *NR* for females and males, respectively. Point colours show the posterior probability of individuals being female (*P(f)*). **Points with black outlines have 95% CI ranges ≤ 0.05 for bootstrapped estimates of *P(f)***. Point shape denotes whether individuals were observed involved in peduncle dives (triangles = receiving, squares = doing, circles = none). The diamond show males observed in the North Atlantic and Arctic Oceans. Dashed vertical lines indicate the minimum body lengths associated with sperm whale sex and age classes defined in **Table 1** as follows: calf (C), juvenile (J), sub-adult (SA), adult female (AF adult male and mature female (AM/MF), maximum female length (Fmax), and mature male (MM).

#### 3.3.2 Posterior probabilities of being female

Models fit with *NR_flipper_* consistently—defined here as having bootstrapped 95% *CI* widths for *P(f)* < 0.05—assigned a high probability (*P(f) > 0.95*) of being female to 21 individuals ranging from 9.8 – 12.5 m *TL* and between 0.28 – 0.31 *NR_flipper_* (**Figure 6)**. This size range slightly exceeds the maximum recorded female length, 12 m ^41^. The *NR_flipper_* model also resulted in a consistently low probability (*P(f)* < 0.05) of individuals being female for 5 individuals between 12.9 – 16.1 m and *NR_flipper_*0.38 – 0.41, which can be classified as males based on their length considerably exceeding typical recorded female lengths (>12).

Both individuals from the North Atlantic fell within the size range expected for males (> 12.5 m) and had relatively large *NR_flipper_* measurements (>0.38), and *P(f)* estimates <0.001 (**Figure 6**).

*P(f)* estimates were robust to varying *chm* and prior *P(f)* values for most individuals, particularly for those that had consistently either high or low probabilities of being female (**Supplementary Material 4**). Individuals for which varying parameter values had a more considerable effect (i.e., > 0.05 difference in *P(f)* between scenarios) had generally intermediate posterior *P(f)* estimates (0.25-0.80), and wide bootstrapped 95% confidence intervals. Images of a sample of individuals and their corresponding *P(f)* values are shown in **Supplementary Material 5**.

#### 3.3.3 Peduncle dive patterns

We inspected 72 recordings (5 – 12 min long) of the footage from which we extracted whale measurements. Within this footage, we found three individuals doing and 12 individuals receiving peduncle dives out of the 90 individuals for which we had at least one total length measurement (**Figure 7**). We were able to measure more individuals receiving peduncle dives than those performing them because the frequent diving involved in performing peduncle dives often resulted in an arched body position which was not suitable for accurate length measurements.

**Figure 7.**
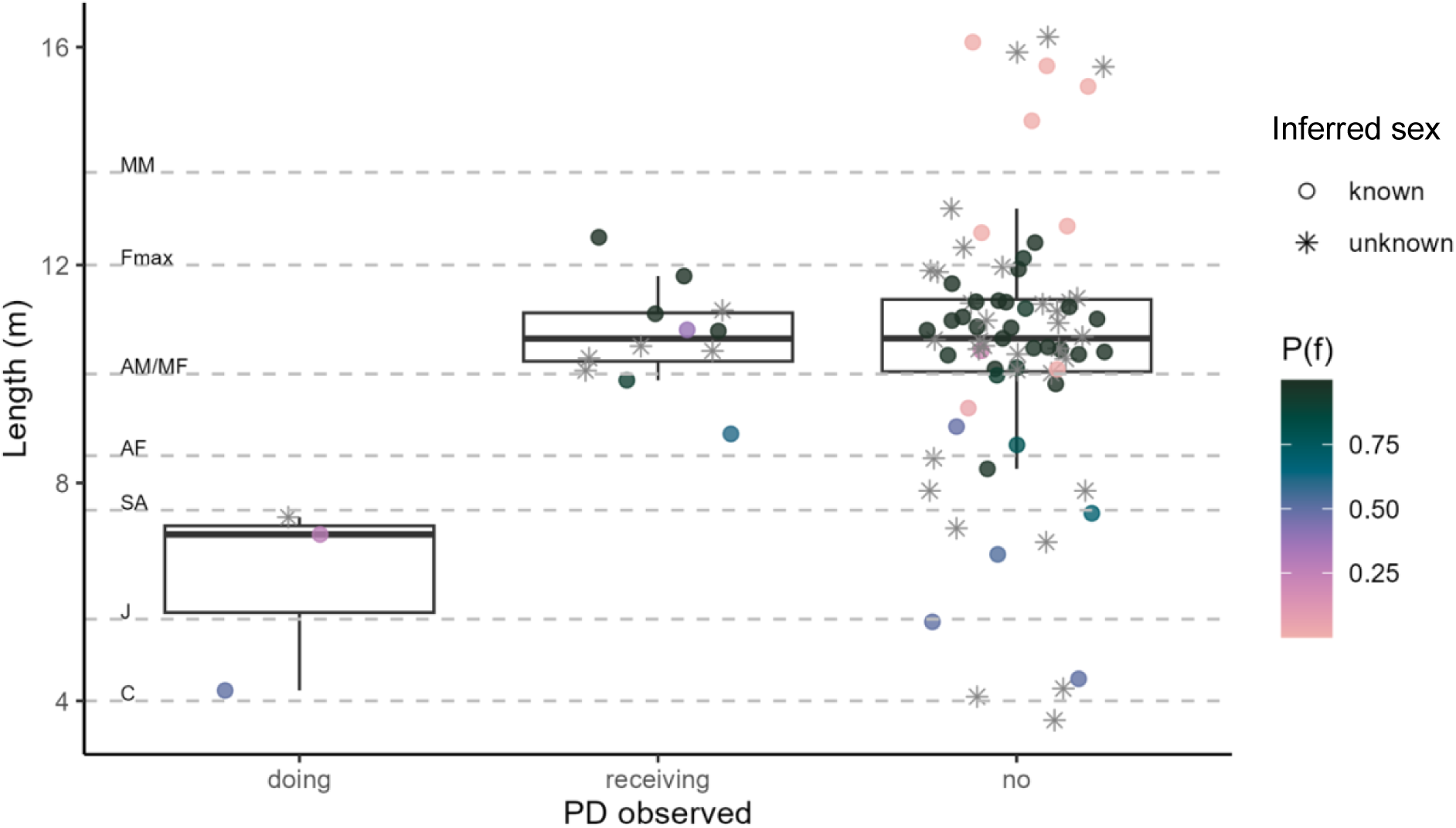
Mean total length (m) distribution of individual sperm whales observed doing, receiving, or not involved in peduncle dives (PD). Points are colored by the bootstrapped mean P(f) estimated using *NR_flipper_* models or are shown as asterisks if *NR_flipper_* could not be measured. Dashed horizontal lines show age-developmental stage classes defined **in Table 1** as follows: calf (C), juvenile (J), sub-adult (SA), adult female (AF adult male and mature female (AM/MF), maximum female length (Fmax), and mature male (MM).

Length measurements of individuals that performed peduncle dives either fell within the total length ranges corresponding to calves (n = 1) or juveniles (n = 3; **Figure 6**). Individuals that received peduncle dives ranged from 8.9 – 12.5 m length, corresponding to the overlapping age/sex classes that include adult to mature females and subadult – adult males. Four out of seven individuals for which we could measure *NR_flipper_* –and thus estimate their probability of being female—had high probability and certainty of being female (*P(f) =* 1, 95% CI [0.99, 1.00]. The remaining three individuals had lower probabilities of being female associated with a high degree of uncertainty (ID11 *P(f)* = 0.38, 95% CI [<0.01, 1.00]; ID75 *P(f)* = 0.61, 95% CI [0.12, 0.96]; ID76 *P(f)* = 0.91, 95% CI [0.52, 1.00]; **Figure 7**).

## 4. DISCUSSION

We developed a minimally invasive method of inferring sperm whale developmental stage and sex by leveraging prior knowledge on sperm whale morphometric development and sexual dimorphism. UAV-based body length (*TL*) estimates provide useful proxies for developmental stages and can help refine the traditionally used ‘calf/mature female-immature/mature male’ classification system. Applying Bayesian theory, we estimated the posterior probabilities of individuals belonging to either sex given their *TL* and *NR_flipper_*. Despite uncertainty arising from different sources of measurement error, we found that nose-to-body ratio measurements based on snout to flipper distances (*NR_flipper_*) reliably captured the development of sexual dimorphism in sperm whales’ noses, with male and female growth curves diverging with increasing length^22,23^. Some individuals could be classified as males or females with high confidence based on their posterior probability estimates, while others lacked the certainty to be assigned as either. Our inspection of peduncle dive patterns (PD) illustrates how our numeric representation of morphological ‘femaleness’ and developmental stage inferences can inform behavioural analyses. Based on simple photogrammetric measurements and a low-cost UAV system, our approach can add key demographic information into sperm whale behavioural analyses and population models.

### 4.1 Developmental stage inferences

Our estimates of total body length were within previously reported size ranges for sperm whales obtained through direct measurements^34,41^. The uncertainty in *TL* estimates of our UAV system (CV = 2.0%) was comparable to most boat-based photogrammetric methods relying on laser photogrammetry (CV = 1.3 - 5.1%^42–44^). However, our UAV system had higher uncertainty than state-of-the-art approaches for measuring sperm whales with UAV systems equipped with laser altimeters (CV = 1.0%^45^). While some research objectives, like detecting individual changes in morphometry over time, may require a higher level of precision, some uncertainty may be acceptable in studies looking at general patterns across a population (e.g. ^46^). Additionally, information on measurement error can be incorporated in statistical analyses, allowing for a measured interpretation of resulting patterns and parameter estimates^10^.

The size-based developmental stage classes we propose refine the existing field-based classification typically used for sperm whales across the globe. The size ranges of our proposed developmental stages (**Table 1**) are based on the size distributions at given developmental milestones (e.g., most individuals that rely exclusively on milk are under 5.5 m long; individuals that incorporate solid foods but still primarily rely on milk (i.e., juveniles) are between 5.5 – 7.6 m long) that are well grounded on anatomical, dietary, and gonadal analyses^34^. Inferences of age or developmental stages based on body size have been used in the past to model population parameters^46^. These inferences generally rely on growth curves that relate individual length measurements to age estimates based on dentin layer counts of killed or stranded individuals^34,47^. However, it is important to note that growth curves are accompanied by uncertainty arising from individual variation in size and development with age^48^. For example, observed *TL* measures for sperm whales have a standard deviation of up to 0.96 m at a given age^46^.

Recent work attempting to identify age-classes based on UAV-derived morphometric measures in common bottlenose dolphins, *Tursiops truncatus,* has shown that size-based age-class assignments perform poorly when age-bin definitions are narrow^13,49^. This highlights the inherent difficulty of converting a continuous measure (*TL*) into a categorical one (age-class). But, when classification bins used for bottlenose dolphins were wide enough, they performed well (with 2 – 3 age class bins having >72.5% individuals assigned within 2 years of actual age-class)^49,50^. Moreover, they found that size-based age classification was most accurate for individuals < 2 years old, which corresponds to the exponential phase of their growth curve, and less accurate once decreasing and stabilizing growth rates are achieved^49^. For sperm whales, the initial exponential growth phase takes place between 0 – 7 years (4.1 – 7.6 m), which would allow for an accurate distinction between calves (<5.5 m) and juveniles (5.5 – 7.6 m).

Additionally, males experience a ‘secondary growth spurt’ after attaining sexual maturity (>10 m), which would make adult males (10 – 13.7 m) reliably distinguishable from mature males (> 13.7 m).

### 4.2 Sex inferences

The overall shape of *NR_flipper_ – TL* growth curves and the resulting posterior probability estimates of individuals being female were generally consistent with previous knowledge on sperm whale sexual dimorphism^22,23^. This was true despite optimum parameter estimates being sensitive to measurement uncertainty (**Figure 4).** Namely, all whales > 13 m—corresponding to the adult/mature male size range^34^—had consistently low probabilities of being female and whales with low *NR_flipper_* (<0.32) between 8.5 – 12 m—corresponding to the mature female size range—had a consistently high probability of being female^34^. Similarly, smaller individuals (<7.6 m) consistently had *P(f)* ∼ 0.5, which is consistent with the expectation that in immature individuals sexual dimorphism, although present, is harder to detect^23^. Hence, our methods allow reliable sex identification (as adult/mature females) of the majority of individuals within the traditional female-immature age-sex class. Likewise, our approach correctly assigned a *P(f) ∼ 0* to the whales observed in the northern North Atlantic and Arctic Oceans, where the vast majority of individuals are known to be adult and mature males^48^. Although this represents a very small sample size (n = 2), it demonstrates the applicability of our methods for data collected in other regions and using different UAV platforms.

Still, our approach resulted in some individuals having high levels of uncertainty and intermediate (i.e. ∼ 0.5) *P(f)* values, despite having *TL* ranges (>8.5 m) at which sexual dimorphism should be detectable based on direct measurements^23^. The uncertainty in *P(f)* estimates may partly be due to the variability associated with our measurement system, particularly for individuals with wide 95% CI estimates.

Intermediate *P(f)* values may also reflect individual variation in levels of sexual dimorphism in secondary sex traits^51^, which would make distinguishing subadult males from adult and mature females particularly challenging. Unfortunately, *NR-TL* curves in ^23^ are based on mean measurements, so we don’t have a baseline for the naturally occurring variation across individuals. Additionally, there are reports across cetacean species of individuals with partial or full hermaphroditism in their genital organs, which in some cases, is linked to intersex chromosome arrangements^52^. Whether these variations translate to ‘intermediate’ secondary sex traits has not been explored. Still, these findings highlight that some caution should be taken when assuming a direct link between phenotype and chromosome arrangement. In the future, individuals found to have uncertain *P(f)* values could be targeted for genetic sampling.

Notably, we found that very few individuals 7.5 - 12.5m had consistently low *P(f)* values—i.e., were likely males—and that those that did fell below the modelled male *NR_flipper_* curve (**Figure 5**). The absence of individuals with higher *NR_flipper_* ratios within this size range may partly reflect the expected departure of young males from their natal units, with most individuals expected to leave when they attain slightly under 10 m (between 7 – 11 years old; **Table 1**^34,46^). Because our fieldwork was focused on large groups which are generally composed of mature females and immature individuals^33^, it is likely that adult (i.e., sexually mature) males were underrepresented in our sample. Despite this, we conservatively assumed equal prior probabilities of observing each sex. This may have underestimated the probability of being female for some intermediate subjects. When we changed the prior expected sex ratio to 0.79 (the estimated proportion of females in the breeding groups off the Galápagos Islands^53^), the posterior probability of individuals with originally intermediate *P(f)* values increased considerably (**Supplement 4 – Figure S4-2)**. However, implementing this informed prior resulted in unrealistically high probabilities of individuals being female for calves (original = 0.5, updated = 0.79), given the even sex ratio of sperm whales in birth and early years^34^. Thus, we consider our conservative prior to produce a better representation of the morphometric difference between males and females throughout their development. In the future, including known sex individuals, especially immature males, may help develop models that can better distinguish the sex of individuals within this size range.

We found consistent support for a constant (i.e., linear) increase of *NR_flipper_* with respect to *TL* for males between 6 – 16.1 m (**Figure 5**). This linear trend in *NR_flipper_* growth emerged despite our initial implementation of a logistic model. The observed pattern aligns with the nose-to-body ratio relationship with body length reported by Nishiwaki et al.^23^.Although our dataset did not cover the full length span of mature males, which can reach over > 18 m^54^, our findings support a continued growth of the male sperm whales’ nose relative to their length despite their net body length growth rate tapering off at 15 m (40 years). This pattern may indicate that the growth in larger males, rather than indeterminate skeletal growth, is primarily driven by the growth of soft tissues encasing the sperm whales’ spermaceti organ, which in older males visibly protrudes beyond the lower jaw when seen from the side^22^.

Sustained growth of secondary sexual traits well beyond sexual maturity has also been observed in other mammal species with high degrees of sexual dimorphism, including giraffes and elephants^55,56^. There is direct evidence that male giraffes with longer necks and larger-bodied elephants have higher reproductive success^55,56^. While the contribution of larger noses to male sperm whales’ reproductive success remains untested, our findings further suggest that strong sexual selective pressures are acting on this trait as it continues to grow despite the potentially high energetic cost of building lipid-rich tissue^22^.

### 4.3 Peduncle dive patterns

Our inspection of PD patterns in relation to inferences of sperm whale age and sex illustrates the applicability of our methods for investigating behavioural patterns. Most (4 of 7) individuals that received PD had consistently high probabilities of being female and ranged between 9.8 – 12.5 m (**Figure 5**), which suggests these are most likely mature females^34^. One individual receiving PD within this size range had relatively low probability of being female, but this estimate was associated with very low certainty (*P(f)* = 0.36, 95% CI width = 1), so its sex and developmental stage are unknown. The remaining individuals observed receiving PDs, for which *P(f)* could not be estimated (n = 5), also fell within the ‘mature female’ size range (**Figure 6**). Two of the individuals observed receiving PDs fell in the ‘adult female’ size range (8.5 – 10 m – **Table 1**) and had high *P(f)* estimates (>0.6), albeit with a low associated certainty (95% CI width > 0.80). While young females of this size range are capable of conceiving^20^, receiving PDs is not exclusive to parous females^57^. Although our approach did not have the resolution to confidently assign all individuals receiving PDs as females, our results generally agree with previous work showing that only females receive peduncle dives^36,37,57^.

We also found all individuals performing peduncle dives were under 7.6 m (**Figure 6**), corresponding to the size range of juveniles (n = 2) and calves (n = 1), which is also congruent with previous work^36,37,57^. Our methods for detecting participation of PD were not exhaustive, as we only inspected a subset of available footage, and thus cannot rule out the participation of any of the remaining individuals in this behaviour. Still, our findings generally aligned with the expectation that this behaviour is limited to calves/juveniles performing the dives, and females receiving them, even if a direct association with suckling remains unclear^37,57^.

### 4.4 Applications

Refined definitions of developmental stages can contribute to our understanding of behavioural development. For instance, investigating the interactions and spatial arrangement between immature individuals and their mothers or caregivers can provide insights into the behavioural development of immature individuals and corresponding changes in maternal care^58–60^. Until now, the systematic study of these behavioural changes has been mostly limited to research on captive individuals or wild populations with extraordinary conditions that allow for longitudinal research approaches (i.e., repeated observations over time of few individuals) to individual behaviour^58,60–62^. Using UAV-derived *TL* estimates, either as continuous or categorical proxies for development, could yield similar insights through a cross-sectional (i.e., observations at a given time across several individuals) approach. This method would be particularly valuable in cases where long-term monitoring and age-determination is impractical, as is the case for highly mobile populations found far offshore. Cooperative care of the young is a central feature and driver of sperm whale sociality^63,64^. Being able to infer the developmental stage and sex of individuals from UAV-derived footage would allow us to better understand the extent to which care behaviours are driven by calves or juveniles seeking care versus adults providing care, and how these change over time. It would also reveal the degree to which each sex may contribute to different aspects of calf care.

Likewise, *TL* and *NR_flipper_* measurements can provide valuable information for interpreting the interactions between adult or mature males and groups of females. It is hypothesized that only mature males (> 13.6 m) participate significantly in reproduction, and that larger males with relatively larger noses have a competitive reproductive advantage^22^, however this has not been empirically tested. By analyzing the interactions between adult/mature females with known males of different sizes and nose-to-body ratio, we would be able to explore if *TL, NR_flipper_*, and raw nose measurements correlate with the frequency with which females approach or interact with males and vice versa. While this would not directly measure reproductive success, patterns of female-male interactions could clarify the drivers of female choice^21^.

Length-based inferences of developmental stage and sex obtained through UAV photogrammetry can also provide a relatively inexpensive and quick method for quantifying the age structure of a population, and inferring its reproductive potential^46,50^. Usually, estimating the age distribution of a population requires large-scale sampling (e.g., hunting or commercial harvesting), mark-recapture methods or long-term monitoring. But, photogrammetric estimates of size distribution, informed by ground-truthing data, can provide useful estimates^46,50^. This is a particularly useful means of monitoring the reproductive potential of a population over time, for instance by comparing reproductive parameters obtained near the end of whaling by traditional methods with present day parameters, which can inform our assessments of populations’ vulnerability with changing conditions. Updating reproductive parameters for sperm whales would contribute to existing knowledge gaps in the different populations’ vulnerability in the face of compounding anthropogenic threats^65^. Still, some care should be taken to make sure that individuals measured are a representative and unbiased sample of the population.

Our methods produce a quantitative representation of the likelihood that an individual is either male or female, which contributes essential information for interpreting behavioural observations. Because differences in the needs between males and females shape their behaviours and dictate their social relationships, the social interactions of males and females can be quite different, especially in sexually dimorphic species. Thus, behavioural studies of social interactions (e.g., affiliative/aversive behaviours, decision-making, cooperation) have been most informative when individual sex can be distinguished (e.g. ^66–68^). This added layer of knowledge can help us make more useful inferences when investigating social interactions. For example, are there social behaviours that are exclusive or predominantly engaged in by mature females? Are some behaviours more frequent among immature males? These questions help elucidate the nature of relationships in sperm whales and the proximate mechanisms by which their societies are maintained and established^62^.

### 4.5 Limitations and methodological considerations

Our work is chiefly limited by the absence of known data on the developmental stage and sex of measured individuals. This means that we cannot provide evaluations of classification performance equivalent to those presented by ^12,13,15^. Future applications of our methods could overcome this limitation by collecting measurements from individuals of known sex and developmental stage. For the present study, we evaluated the ability of our methods to infer individual developmental stages and sexes by comparing our findings to those based on direct measurements of thousands of killed individuals^23,34,47^ or mass strandings^41^. While these sources provide a useful baseline, there are some caveats to extrapolating these findings to our sample. Beyond individual variation in growth rates, population-level growth rates can change in response to resource availability and human impacts^69^. For instance, ^70^ found that female sperm whales killed before the whaling moratorium (1959 – 1962) in the Eastern South Pacific sexually matured earlier (6.5 years) and at smaller sizes (8.2 m) than in other regions, presumably as a result of prolonged whaling in the region. Similarly, ^46^ found that growth curves and overall lengths of Galápagos sperm whales in 1985 and 1987 were slightly smaller than those generated in previous decades using whaling data. While some of the differences in the latter case may reflect a bias in whaling data towards larger and more lucrative individuals in whaling data, the differences between growth curves remained within the expected variation of age – length relationships^46^. There is also evidence that size distributions among female sperm whales vary geographically, with whales in lower latitudes being generally smaller than those in higher latitudes^71^. Thus, while our general appraisal of developmental stage and sex is informative, the precise parameters describing the *TL* and *NR_flipper_* curves may not be directly applicable to whales from other regions. Applying this method to other datasets will require estimating optimal parameters for a given population.

We chose an UAV system that is relatively inexpensive (<500 USD vs > 2,000 USD for other frequently used systems) and user-friendly, which may be ideal for projects that are budget and/or experience- limited, allowing them to collect valuable demographic data that would otherwise not be attainable. If higher accuracy and precision are needed, simple improvements can be made by implementing laser-based altimeters. There are several open-sourced resources for installing lidar systems on commercially available UAVs frequently used in cetacean monitoring^39^. This would improve accuracy and precision in *TL* estimates^10,72^, but would not resolve the uncertainty associated with measuring *NR_flipper_* (or other body ratios) as they are independent of altitude estimates. Because whales are not rigid, there is some unavoidable uncertainty in this. We suggest, as we have done here, taking several measures for the same individuals to quantify the uncertainty associated with these metrics, which can then be propagated through further analytical steps using frequentist or Bayesian approaches (e.g. ^10,72^).

## 5 Conclusions

Our work demonstrates the application of UAV-based photogrammetry to infer the developmental stage and sex of live sperm whales. By combining modern techniques with historical whaling data, we developed a minimally invasive, low-cost means of obtaining refined demographic data that represents a significant advancement to traditional age-class assignments that were possible based on field observations. In the future, this approach can be further refined by incorporating measures of individuals with known sex/age and adoption of state-of-the art UAV systems. The ability to obtain individual sex and developmental stage inferences will contribute to our understanding of the development and divergence of social behaviour of this extremely sexually dimorphic species. Moreover, finer-scale demographic data provides key information for evaluating populations and reproductive potential, which is be particularly valuable in regions with unfavourable or uncertain conservation status. Finally, while our approach to inferring sex was developed based on sperm whales’ particular morphological development, it may be adapted towards other species with morphological differences between sexes emerging before full maturity.

## Acknowledgements

We thank the crew and skippers that participated in the 2023 Galápagos field season, particularly Luke Rendell, Hansen Johnson, and Mateo Valencia. We are also grateful to the Galápagos National Park Service for granting authorization to conduct research in the region (Research permit No. PC-86-22) and the Charles Darwin Foundation, especially Marta Romoleroux, for their invaluable support of our operations. Fieldwork was possible through funding from the Natural Sciences and Engineering Research Council of Canada (NSERC) and a Rufford Small Grant. AE was supported through the Killam Trust; CC and HW were funded through NSERC.

## References

1. Bleich, V. C., Bowyer, R. T. & Wehausen, J. D. Sexual segregation in mountain sheep: resources or predation? Wildl. Monogr. 134, 3–50 (1997).

2. Ruckstuhl, K. E. Sexual segregation in vertebrates: proximate and ultimate causes. Integr. Comp. Biol. 47, 245–257 (2007).

3. Griffiths, S. W., Orpwood, J. E., Ojanguren, A. F., Armstrong, J. D. & Magurran, A. E. Sexual segregation in monomorphic minnows. Anim. Behav. 88, 7–12 (2014).

4. Volis, S. & Deng, T. Importance of a single population demographic census as a first step of threatened species conservation planning. Biodivers. Conserv. 29, 527–543 (2020).

5. Le Clercq, L., Kotzé, A., Grobler, J. P. & Dalton, D. L. Biological clocks as age estimation markers in animals: a systematic review and meta-analysis. Biol. Rev. 98, 1972–2011 (2023).

6. Shaw, C. N., Wilson, P. J. & White, B. N. A reliable molecular method of gender determination for mammals. J. Mammal. 84, 123–128 (2003).

7. Gowans, S., Whitehead, H. & Hooker, S. K. Social organization in northern bottlenose whales, *Hyperoodon ampullatus*: not driven by deep-water foraging? Anim. Behav. 62, 369–377 (2001).

8. Denkinger, J. et al. Social structure of killer whales (*Orcinus orca*) in a variable low-latitude environment, the Galápagos Archipelago. Mar. Mammal Sci. mms.12672 (2020) doi:10.1111/mms.12672.

9. Burnett, J. D. et al. Estimating morphometric attributes of baleen whales with photogrammetry from small UASs: A case study with blue and gray whales. Mar. Mammal Sci. 35, 108–139 (2019).

10. Bierlich, K. et al. Bayesian approach for predicting photogrammetric uncertainty in morphometric measurements derived from drones. Mar. Ecol. Prog. Ser. 673, 193–210 (2021).

11. Glarou, M. et al. Estimating body mass of sperm whales from aerial photographs. Mar. Mammal Sci. 39, mms.12982 (2022).

12. Vivier, F. et al. Inferring dolphin population status: using unoccupied aerial systems to quantify age-structure. Anim. Conserv. acv.12978 (2024) doi:10.1111/acv.12978.

13. Cheney, B. J., Dale, J., Thompson, P. M. & Quick, N. J. Spy in the sky: A method to identify pregnant small cetaceans. *Remote Sens*. Ecol. Conserv. 8, 492–505 (2022).

14. Fernandez Ajó, A., et al. Assessment of a non-invasive approach to pregnancy diagnosis in gray whales through drone-based photogrammetry and faecal hormone analysis. R. Soc. Open Sci. 10, (2023).

15. Robinson, C. V. & Visona-Kelly, B. C. A geometric morphometric approach for detecting different reproductive stages of a free-ranging killer whale *Orcinus orca* population. Sci. Rep. 15, (2025).

16. Whitehead, H. Sperm Whales: Social Evolution in the Ocean. (University of Chicago Press, Chicago, USA, 2003).

17. Cantor, M., Eguiguren, A., Merlen, G. & Whitehead, H. Galápagos sperm whales (*Physeter macrocephalus*): Waxing and waning over three decades. Can. J. Zool. 95, 645–652 (2017).

18. Whitehead, H., Coakes, A., Jaquet, N. & Lusseau, S. Movements of sperm whales in the tropical Pacific. Mar. Ecol. Prog. Ser. 361, 291–300 (2008).

19. Christal, J. & Whitehead, H. Aggregations of mature male sperm whales on the Galápagos Islands breeding ground. Mar. Mammal Sci. 13, 59–69 (1997).

20. 20. Rice, D. W. Sperm whale. Physeter macrocephalus Linnaeus, 1758. in Handbook of marine mammals (eds Ridgway, S. H. & Harrison, R.) 177–233 (Academic Press, London, 1989).

21. Eguiguren, A., Konrad Clarke, C. M. & Cantor, M. Sperm whale reproductive strategies: Current knowledge and future directions. in Sex in Cetaceans: Morphology, Behavior, and the Evolution of Sexual Strategies (eds Würsig, B. & Orbach, D. N.) 443–467 (Springer International Publishing, Cham, 2023). doi:10.1007/978-3-031-35651-3_19.

22. Cranford, T. W. The sperm whale’s nose: sexual selection on a grand scale? Mar. Mammal Sci. 15, 1133–1157 (1999).

23. Nishiwaki, M., Ohsumi, S. & Maeda, Y. Change of form in the sperm whale accompanied with growth. Sci. Rep. Whales Res. Inst. Tokyo 17, 1–17 (1963).

24. Altmann, J. Observational study of behavior: sampling methods. Behaviour 49, 227–266 (1974).

25. Napoli, C. et al. Drone-based photogrammetry reveals differences in humpback whale body condition and mass across North Atlantic foraging grounds. Front. Mar. Sci. 11, 1336455 (2024).

26. Torres, W. & Bierlich, K. MorphoMetriX: a photogrammetric measurement GUI for morphometric analysis of megafauna. J. Open Source Softw. 5, 1825 (2020).

27. Friard, O. & Gamba, M. BORIS: A free, versatile open-source event-logging software for video/audio coding and live observations. Methods Ecol. Evol. 7, 1325–1330 (2016).

28. O’Callaghan, S. A. et al. Aerial photo-identification of sperm whales (*Physeter macrocephalus*). Aquat. Mamm. 50, 479–494 (2024).

29. Arnbom, T. Individual identification of sperm whales. Int. Whal. Commision 37, 201–204 (1987).

30. R Core Team. R: A language and environment for statistical computing. R Foundation for Statistical Computing (2019).

31. Dixon, P. M. The bootstrap and the jackknife: Describing the precision of ecological indices. in Design and Analysis of Ecological Experiments (eds Scheiner, S. M. & Gurevitch, J.) 267–288 (Oxford University PressNew York, NY, 2001). doi:10.1093/oso/9780195131871.003.0014.

32. Richard, K. R., Dillon, M. C., Whitehead, H. & Wright, J. M. Patterns of kinship in groups of free-living sperm whales (*Physeter macrocephalus*) revealed by multiple molecular genetic analyses. Proc. Natl. Acad. Sci. 93, 8792–8795 (1996).

33. 33. Best, P. B. Social organization in sperm whales, *Physeter macrocephalus*. in Behavior of marine animals (eds Winn, H. E. & Olla, B. L.) 227–290 (University of California Press, Berkeley, 1979).

34. Best, P. B., Canham, P. A. S. & Macleod, N. Patterns of reproduction in sperm whales, Physeter macrocephalus. Rep. Int. Whal. Comm. 51–79 (1984).

35. Mendes, S., Newton, J., Reid, R. J., Zuur, A. F. & Pierce, G. J. Stable carbon and nitrogen isotope ratio profiling of sperm whale teeth reveals ontogenetic movements and trophic ecology. Oecologia 151, 605–615 (2007).

36. Gero, S. & Whitehead, H. Sucking behavior in sperm whale calves: observations and hypotheses. Mar. Mammal Sci. 23, 398–413 (2007).

37. Sarano, F. et al. Nursing Behavior in Sperm Whales (*Physeter macrocephalus*). Anim. Behav. Cogn. 10, 105–131 (2023).

38. Konrad, C. M., Frasier, T. R., Whitehead, H. & Gero, S. Kin selection and allocare in sperm whales. Behav. Ecol. 30, 194–201 (2019).

39. Bierlich, K. C. et al. LidarBoX: A 3D-printed, open-source altimeter system to improve photogrammetric accuracy for off-the-shelf drones. Drone Syst. Appl. 12, 1–10 (2024).

40. Whitehead, H., Brennan, S. & Grover, D. Distribution and behaviour of male sperm whales on the Scotian Shelf, Canada. Can. J. Zool. 70, 912–918 (1992).

41. Evans, K. & Hindell, M. A. The age structure and growth of female sperm whales (*Physeter macrocephalus*) in southern Australian waters. J. Zool. 263, 237–250 (2004).

42. Gordon, J. A simple photographic technique for measuring the length of whales from boats at sea. Rep. Int. Whal. Comm. 40, 581–588 (1990).

43. Dawson, S. M., Chessum, C. J., Hunt, P. J. & Slooten, E. An inexpensive stereophotographic technique to measure sperm whales from small boats. Rep. Int. Whal. Comm. 45, 431–436 (1995).

44. Jaquet, N. A simple photogrammetric technique to measure sperm whales at sea. Mar. Mammal Sci. 22, 862–879 (2006).

45. Dickson, T., Rayment, W. & Dawson, S. Drone photogrammetry allows refinement of acoustically derived length estimation for male sperm whales. Mar. Mammal Sci. 37, 1150–1158 (2021).

46. Waters, S. & Whitehead, H. Population and growth parameters of Galápagos sperm whales estimated from length distributions. Rep. Int. Whal. Comm. 40, 225–235 (1990).

47. Ohsumi, S. Age-length key of the male sperm whale in the North Pacific and comparison of growth curves. Rep. Int. Whal. Comm. 27, 295–300 (1977).

48. Martin, A. R. An examination of sperm whale age and length data from the 1949-78 Icelandic catch. Rep. Int. Whal. Comm. 30, 227–231 (1980).

49. Vivier, F. et al. Quantifying the age structure of free-ranging delphinid populations: Testing the accuracy of Unoccupied Aerial System photogrammetry. Ecol. Evol. 13, (2023).

50. Vivier, F. et al. Inferring dolphin population status: using unoccupied aerial systems to quantify age-structure. Anim. Conserv. 28, 262–276 (2025).

51. McLaughlin, J. F. et al. Multivariate models of animal sex: breaking binaries leads to a better understanding of ecology and evolution. Integr. Comp. Biol. 63, 891–906 (2023).

52. Einfeldt, A. L., Orbach, D. N. & Feyrer, L. J. A method for determining sex and chromosome copy number: sex-by-sequencing reveals the first two species of marine mammals with XXY chromosome condition. J. Mammal. 100, 1671–1677 (2019).

53. 53. Richards, A. F. Life history and behavior of female dolphins (Tursiops sp.) in Shark Bay, Western Australia. (University of Michigan, United States -- Michigan, 1996).

54. Kasuya, T. Density dependent growth in North Pacific sperm Whales. Mar. Mammal Sci. 7, 230–257 (1991).

55. Simmons, R. E. & Scheepers, L. Winning by a neck: sexual selection in the evolution of giraffe. Am. Nat. 148, 771–786 (1996).

56. Hollister-Smith, J. A. et al. Age, musth and paternity success in wild male African elephants, *Loxodonta africana*. Anim. Behav. 74, 287–296 (2007).

57. Konrad, C. M., Frasier, T. R., Whitehead, H. & Gero, S. Kin selection and allocare in sperm whales. Behav. Ecol. 30, 194–201 (2019).

58. Mann, J. & Smuts, B. B. Natal attraction: allomaternal care and mother–infant separations in wild bottlenose dolphins. Anim. Behav. 55, 1097–1113 (1998).

59. Mann, J. & Smuts, B. Behavioral development in wild bottlenose dolphin newborns (*Tursiops* sp.). Behaviour 136, 529–566 (1999).

60. Fellner, W., Bauer, G. B., Stamper, S. A., Losch, B. A. & Dahood, A. The development of synchronous movement by bottlenose dolphins (*Tursiops truncatus*). Mar. Mammal Sci. 29, E203– E225 (2013).

61. Sakai, M. et al. Mother-calf interactions and social behavior development in Commerson’s dolphins (*Cephalorhhynchus commersonii*). J. Ethol. 31, 305–313 (2013).

62. Eguiguren, A., Walmsley, S. F., Feyrer, L. J., Zwamborn, E. M. J. & Whitehead, H. The role of touch in marine mammal sociality: A review and future directions. Preprint at 10.32942/X2NM0Q (2025).

63. Gero, S., Gordon, J. & Whitehead, H. Calves as social hubs: Dynamics of the social network within sperm whale units. Proc. R. Soc. B Biol. Sci. 280, 20131113 (2013).

64. Cantor, M., Gero, S., Whitehead, H. & Rendell, L. Sperm whale: The largest toothed creature on earth. in Ethology and Behavioral Ecology of Odontocetes (ed. Würsig, B.) 261–280 (Springer International Publishing, Cham, 2019). doi:10.1007/978-3-030-16663-2_12.

65. Eguiguren, A. et al. Integrating cultural dimensions in sperm whale (*Physeter macrocephalus*) conservation: threats, challenges and solutions. Philos. Trans. R. Soc. B Biol. Sci. 380, 20240142 (2025).

66. Connor, R. C., Mann, J. & Watson-Capps, J. A sex-specific affiliative contact behavior in Indian Ocean bottlenose dolphins, *Tursiops* sp. Ethology 112, 631–638 (2006).

67. Harvey, B. S., Dudzinski, K. M. & Kuczaj, S. A. Associations and the role of affiliative, agonistic, and socio-sexual behaviors among common bottlenose dolphins (*Tursiops truncatus*). Behav. Processes 135, 145–156 (2017).

68. Zwamborn, E. M. J., Walmsley, S. F. & Whitehead, H. Flanking female guides: Collective decision making in long-finned pilot whales. Anim. Behav. 205, 149–159 (2023).

69. Adamczak, S. K. et al. Growth in marine mammals: A review of growth patterns, composition and energy investment. Conserv. Physiol. 11, coad035 (2023).

70. Clarke, R., Paliza, O. & Waerebeek, K. V. Sperm whales of the Southeast Pacific. Part VII. Reproduction and growth in the female. 32 (2012).

71. Best, P. B., Tormosov, D., Brandão, A. & Mikhalev, Y. Geographical variation in the body size of adult female sperm whales (*Physeter macrocephalus*) – an example of McNab’s resource rule? Mammalia 81, 189–196 (2016).

72. Napoli, C. et al. Drone-based photogrammetry reveals differences in humpback whale body condition and mass across North Atlantic foraging grounds. Front. Mar. Sci. 11, 1336455 (2024).

73. 73. Norris, K. S. & Harvey, G. W. A theory for the function of the spermaceti organ of the sperm whale (*Physeter catodon L*.). in Animal Orientation and Navigation (eds Galler, S. R., Schmidt-Koenig, K., Jacobs, G. J. & Belleville, R. E.) vol. 262 (NASA Scientific and Technical Office, USA, 1972).

74. Madsen, P. T., Whalberg, M. & Møhl, B. Male sperm whale (*Physeter macrocephalus*) acoustics in a high-latitude habitat: Implications for echolocation and communication. Behav. Ecol. Sociobiol. 53, 31–41 (2002).

75. Clarke, R. & Paliza, O. Intraspecific fighting in sperm whales. Rep. Int. Whal. Comm. 38, 235–241 (1988).

76. Carrier, D. R., Deban, S. M. & Otterstrom, J. The face that sunk the Essex: Potential function of the spermaceti organ in aggression. J. Exp. Biol. 205, 1755–1763 (2002).

77. Panagiotopoulou, O., Spyridis, P., Mehari Abraha, H., Carrier, D. R. & Pataky, T. C. Architecture of the sperm whale forehead facilitates ramming combat. PeerJ 4, e1895 (2016).

78. Gero, S. et al. Behavior and social structure of the sperm whales of Dominica, West Indies. Mar. Mammal Sci. 30, 905–922 (2014).

