## Supplementary Material for "Inferring sperm whale (*Physeter macrocephalus*) sex and developmental stage using aerial photogrammetry" (from another version of this preprint: version 1, 2025-10-17)

**Supplementary Material 1**

*-Error estimation for DJI Mini Drone 2-*

### S1.1 Altitude error estimation and correction

We ran three candidate hierarchical generalized linear models to correct drone altitude using the *nlme* R package (Pinheiro et al. 2020)

1. True altitude ~ barometric altitude (no random effects)
2. True altitude ~ barometric altitude (random intercept = Date)
3. True altitude ~ barometric altitude (random intercept & slope = Date)

Adding date as a random effect was intended to capture the effect that different weather conditions could have on barometric altitude measurements. We found that models including date as a random variable improved overall model performance, as demonstrated by the significantly lower AIC for models 2 and 3 (**Table S1 1**). Model 3 also resulted in corrected altitude measurements with narrower confidence. Still, hierarchical models decreased measurement error and uncertainty marginally, with model 3 resulting in 95% CI error width being only 20 cm smaller than model 1. While this difference may be important in some contexts (e.g., when attempting to detect growth or changes in body condition for individuals), we considered it negligible to our goal of inferring general developmental stages and sex.

***Table S1 1.*** Summary statistics (mean, standard deviation (SD), 2.5$^{th}$ percentile, 97.5$^{th}$ percentile, and 95% CI width) for the error (m) estimates of drone altitude and calibration object length (of a research vessel with true length 12.03) based on the corrected drone altitude. dAIC shows the difference in AIC from the model with the lowest AIC. Raw, uncorrected altitude values are shown in the first row. The final model (1) used in this study is highlighted in gray.

| | | **Altitude error (m)** | | | | | **Measurement error (m)** | | | | |
|---|---|---|---|---|---|---|---|---|---|---|---|
| **Correction model** | ΔAIC | Mean | SD | 2.5$^{th}$ p | 97.5$^{th}$ p | 95% CI width | Mean | SD | 2.5$^{th}$ p | 97.5$^{th}$ p | 95% CI width |
| uncorrected | NA | 2.4 | 1.9 | -1.2 | 0.2 | 1.4 | -0.55 | 0.38 | -1.23 | 0.16 | 1.39 |
| 1. No random effects | 65.6 | 0.00 | 1.9 | -3.9 | -2.1 | 1.8 | 0.02 | 0.38 | -0.65 | 0.74 | 1.39 |
| 2. Date as random **intercept** | 50.2 | 0.00 | 1.8 | -3.8 | -1.8 | 1.9 | 0.01 | 0.36 | -0.69 | 0.70 | 1.38 |
| 3. Date as random **intercept** and **slope** | 0 | 0.00 | 1.5 | -2.7 | -1.7 | 1.0 | 0.01 | 0.33 | -0.61 | 0.59 | 1.20 |

### Supplementary Material 2

*-UAV footage quality rating scale-*

Each video was rated based on the sum of the score assigned to each aspect described in **Table S2-1.** Only video recordings with a score <5 were used to extract sperm whale measurements. Although conditions sometimes varied within a video recording, the score given to each aspect was intended to capture most of the footage.

***Table S2-1.*** Definition and score levels of aspects used to measure video footage quality.

| **Aspect** | **Definition** | **0** | **1** | **2** |
|---|---|---|---|---|
| Glare | The reflection or shine on the surface of the water produced by the sun | Glare covers <25% of the frame or is absent | Glare of medium intensity covers > 25% of the frame | Glare of high intensity covers > 25% of the frame |
| Sea state | The texture of the water which can interact with glare or distort the whale's body. (Not to be confused with Beaufort scale) | Flat calm. Swell or undulations may be visible, but no ripples or waves. | Small ripples are visible. | Large ripples or waves are visible, distorting the surface of the water. |
| Focus | The sharpness with which individuals can be observed in the water. | Edges are sharp and details (like body parts) are easily observed on whales. | Edges are slightly fuzzy and only some details are visible. | Edges are fuzzy and details can't be discerned. |
| Exposure | Level of light that is captured in an image. | Level of light is adequate, and the video does not require adjustments to be able to observe individuals | Image is slightly dark or light, but it can be fixed by modifying image setting digitally | Image is too dark or light and can't be fixed digitally. |

**Supplementary Material 3**

*-Results summary for sex inferences based NRdorsal measurements in comparison with NRflipper measurements-*

### S3.1 Variability in nose-to-body ratio measurements

Measurements of $NR_{dorsal}$ were generally more variable than measurements of $NR_{flipper}$, with nearly twice as many individuals having wider 95% CI's for $NR_{dorsal}$ measurements (n = 33) than $NR_{flipper}$ measurements (n = 18; **Figure S3-1**). This likely reflects the absence of a reliably distinct boundary for the base of the dorsal fin in several whales.

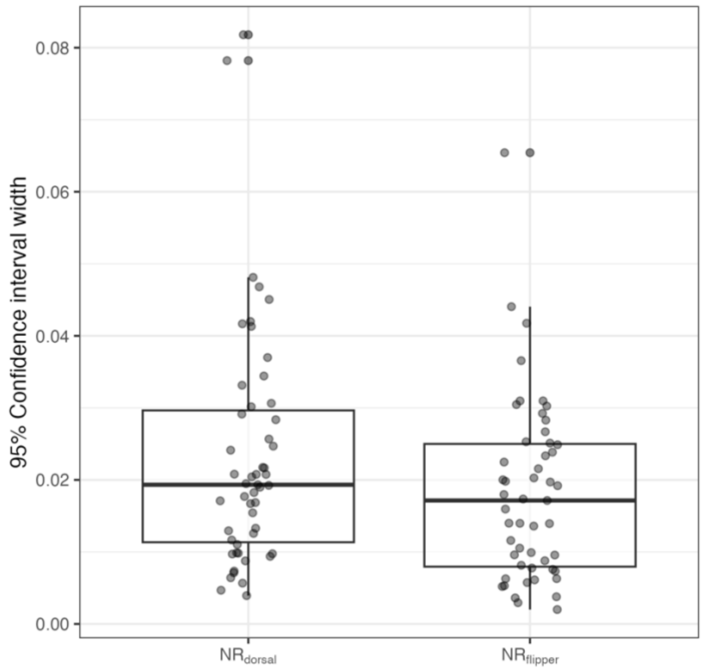


***Figure S3-1.*** Distribution of 95% Confidence interval widths for bootstrapped $NR_{dorsal}$ and $NR_{flipper}$ measurements.

### S3.2 Parameter optimization

Optimal values for *fr* and $m_r$ varied more across bootstrap iterations of the models fit with $NR_{dorsal}$ than $NR_{flipper}$ (**Table S3-1**), resulting in generally higher levels of uncertainty associated with models based on $NR_{dorsal}$ than $NR_{flipper}$ (**Figure S3-2** & **Figure S3-3**).

***Table S3-1*** Bootstrapped means and 95$^{th}$ percentile confidence intervals (95% CI) based on 1000 iterations for parameters relating sperm whale length (m) and nose-to-body ratio (NR) metrics based on snout to the caudal base of the dorsal fin ($NR_{dorsal}$) and on snout to the flipper insertion point ($NR_{flipper}$). Parameters reflect the growth rate of females and small males (≤ 6 m) (fr), the female asymptote of R ($max_f$), the growth rate of larger males (> 6 m) (mr), and the male asymptote of R ($max_m$).

| ***R* Metric** | ***fr* [95% CI]** | **$max_f$ [95% CI]** | ***mr* [95% CI]** | **$max_m$ [95% CI]** |
|---|---|---|---|---|
| $NR_{dorsal}$ | 1.28 (0.66 - 2.38) | 0.65 (0.64 - 0.65) | 0.17 (0.05 - 0.4) | 0.36 (0.22 - 0.72) |
| $NR_{flipper}$ | 0.72 (0.50 - 1.11) | 0.3 (0.3 - 0.3) | 0.07 (0.02 - 0.15) | 0.81 (0.48 - 1.73) |

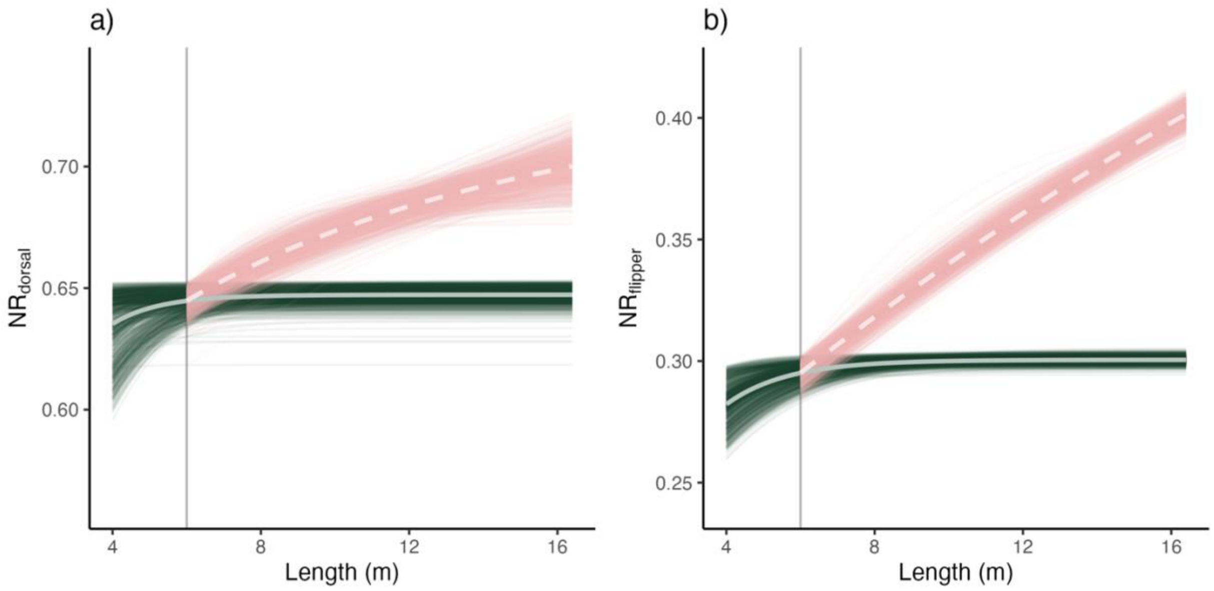


***Figure S3-2*** Bootstrapped logistic curves of the total length (m) and the nose-to-body ratio of sperm whales based on measures of the snout to the caudal base of the dorsal fin (a) and snout to the base of the flipper (b). Theoretical male curves are shown in violet and theoretical female curves are shown in green. The average NR values across iterations are shown by light violet dashed and green solid lines for males and females, respectively. The vertical line indicates the point of divergence between males and females (chm = 6 m) based on (Nishiwaki et al. 1963)

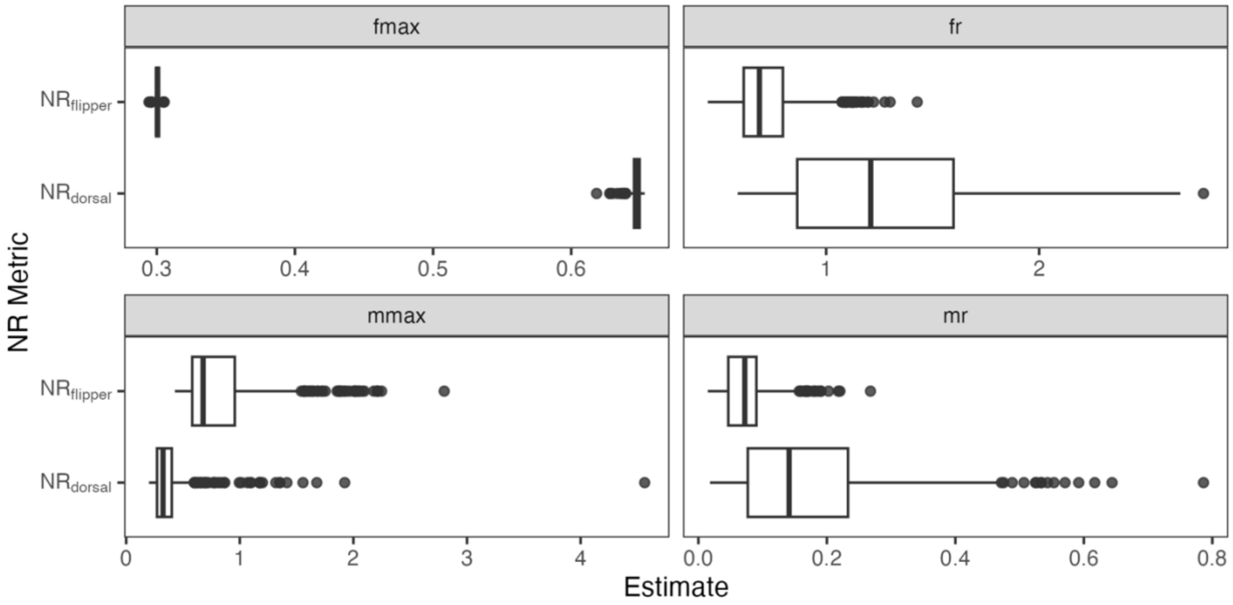


***Figure S3-3*** Distribution of bootstrapped parameter estimates (x axis) for $NR_{dorsal}$ and $NR_{flipper}$ models.

#### S3.3 Posterior probabilities of being female

Models based on $NR_{flipper}$ metrics resulted in more reasonable individual estimates of the probability of being female than $NR_{dorsal}$ based models. For example, individual ID74 (mean $TL$ = 10.78 m, 95% $CI$ = 10.63 – 11.06 m), which was observed receiving peduncle dives, was classified with high confidence as female by $NR_{flipper}$ models (mean $P(f)$ = 0.99, 95% $CI$ = 0.99 – 1.00), yet received low and uncertain estimates from the $NR_{dorsal}$ model (mean = 0.12, 95% $CI$ = 0 – 0.44; **Figure S3-4**). Similarly, individual ID04, a large male (mean $TL$ = 15.2 m, 95% $CI$ = 14.9 – 15.5 m), was confidently assigned a near-zero probability of being female by the $NR_{flipper}$ model (mean < 0.001, 95% *CI width* = 0), but received an uncertain and intermediate probability estimate based on the $NR_{dorsal}$ models (mean = 0.50, 95% $CI$ = <0.001 – 0.97). Additionally, in models fit with $NR_{dorsal}$, only two individuals that could be assumed to be mature males based on their sizes (ID01 & ID81) were consistently assigned low probabilities of being females. No individuals were consistently assigned a high probability of being female based on $NR_{dorsal}$ models.

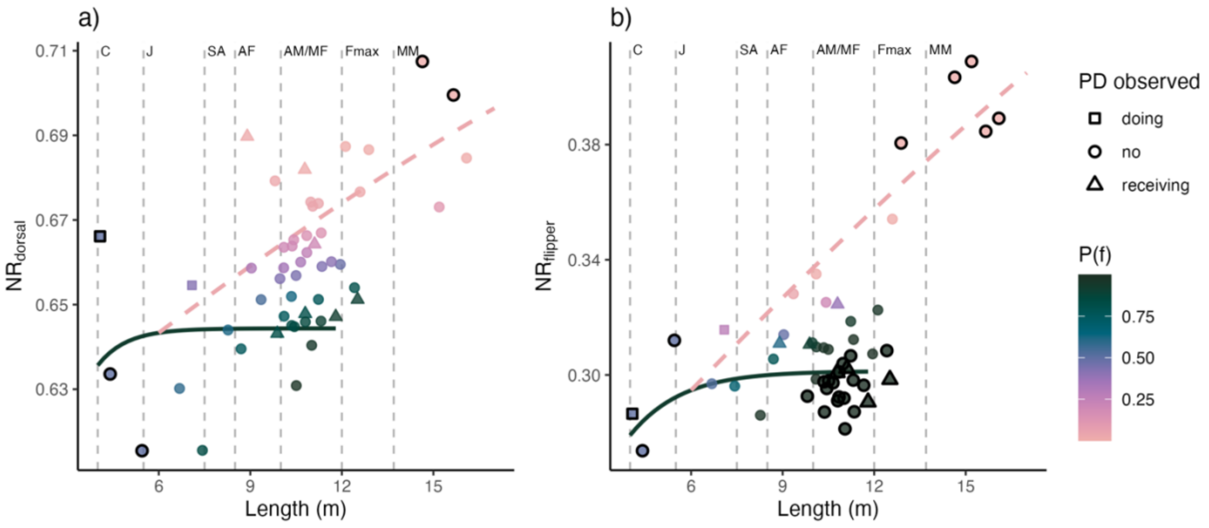


***Figure S3-4*** Bootstrapped mean Total Length (m) and nose-to-body ratio (NR) for individual sperm whales based on (a) snout – dorsal fin distance ($NR_{dorsal}$) and (b) snout – flipper distance ($NR_{flipper}$). The solid green line and dashed pink lines show the bootstrapped mean modeled $NR$ for females and males, respectively. Point colours show the mean posterior probability of individuals being female (P(f)). **Points with black outlines have 95% CI ranges ≤ 0.05 for bootstrapped estimates of *P(f)*.** Point shape denotes whether individuals were observed involved in peduncle dives (triangles = receiving, squares = doing, circles = none). Dashed vertical lines indicate the minimum body lengths associated with sperm whale sex and age classes as follows: calf (4 m; NB), juvenile (J; 5.5 m), sub-adult (SA; 7.6 m ), adult female (AF – 8.5 m), adult male and mature female (AM/MF – 10 m), maximum female length (Fmax – 12 m), and mature male (MM – 13.7).

### Supplementary Material 4

*-Robustness checks-*

#### S4.1 Effect of varying *chm* values on posterior *P(f)* estimates

Varying *chm* values between 5 – 7 m resulted in little change in *P(f)* estimates for most individuals (41 of 50), with estimates differing by less than 0.05 across *chm* values (**Figure S4-1**). The individuals for which varying *chm* values had more considerable effects also had intermediate *P(f)* estimates (0.25-0.80), and wide bootstrapped 95% confidence intervals (> 0.5),

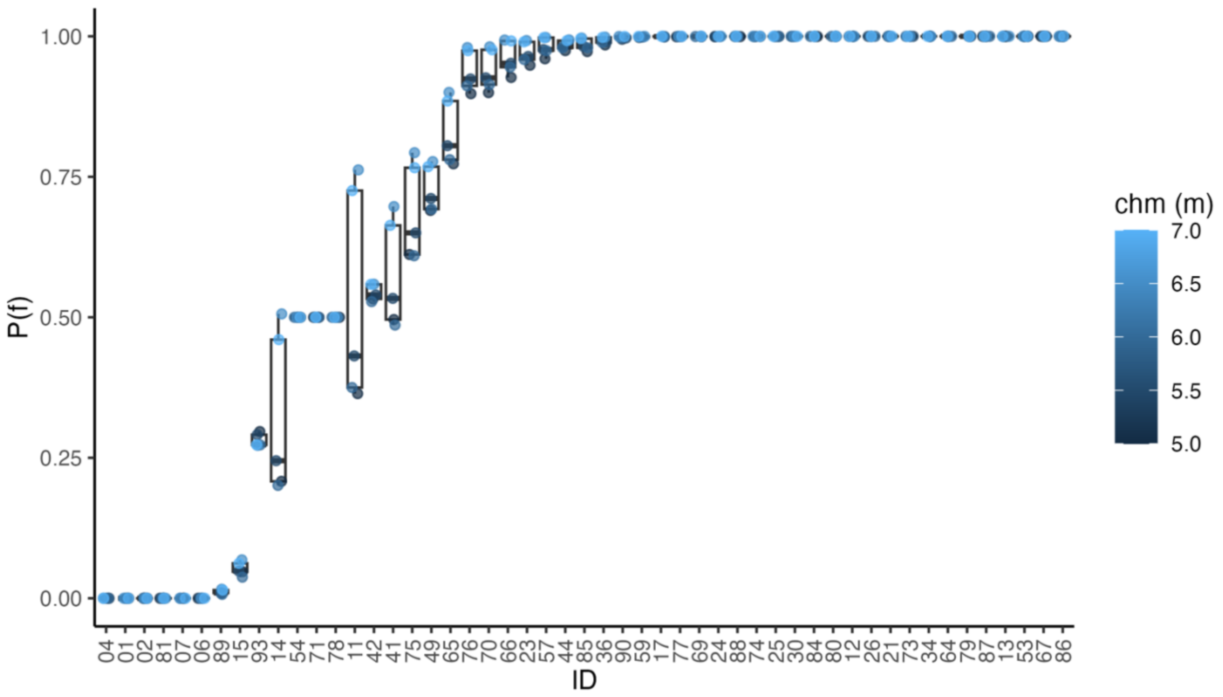


***Figure S4-1.*** Probabilities of individuals (ID) being female (*P(f)*) resulting from varying the length at divergence (*chm*) in male and female nose-to-body ratio growth curves. Individuals are sorted by average *P(f)*. Points show individual estimates coloured by corresponding *chm* values.

#### S4.2 Effect of varying prior probabilities on posterior *P(f)* estimates

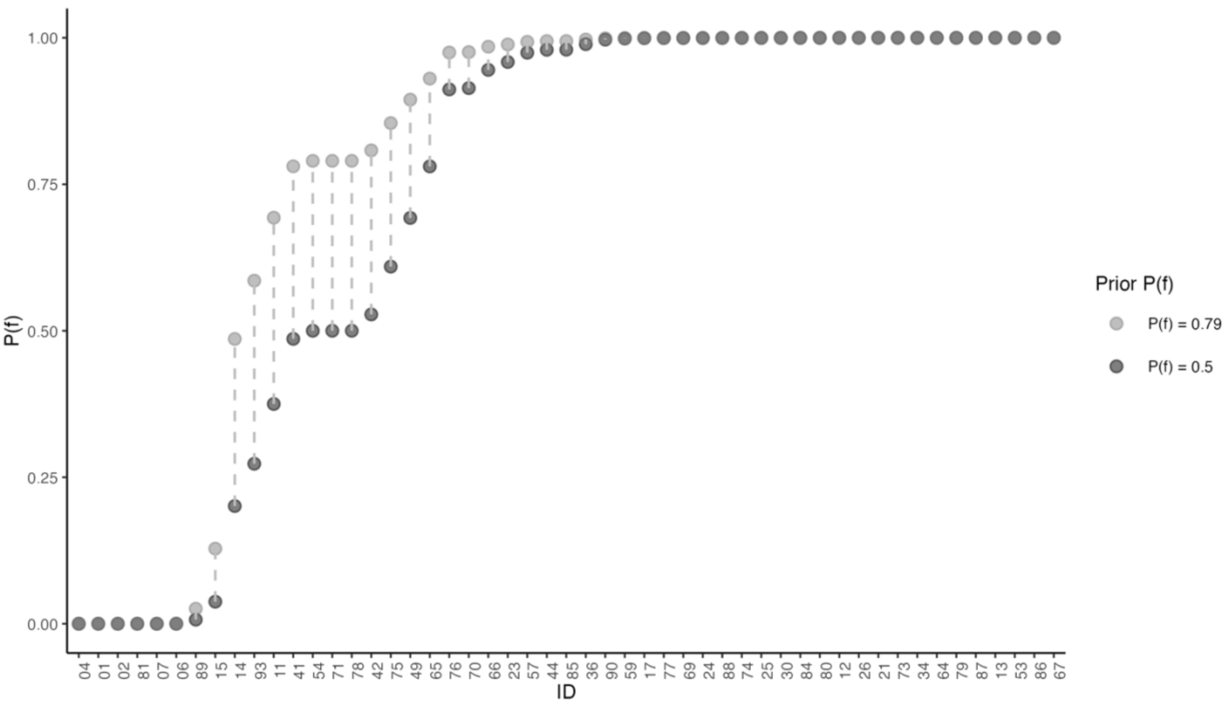


***Figure S4-2.*** Probabilities of individuals (ID) being female (*P(f)*) based on conservative prior assumptions (dark gray, prior *P(f)* = 0.5) and informed prior assumptions (light gray, *P(f)* = 0.79). Individuals are sorted by *P(f)* estimates based on conservative prior assumptions.

### Supplementary Material 5

*-Sample photographs of measured sperm whales-*

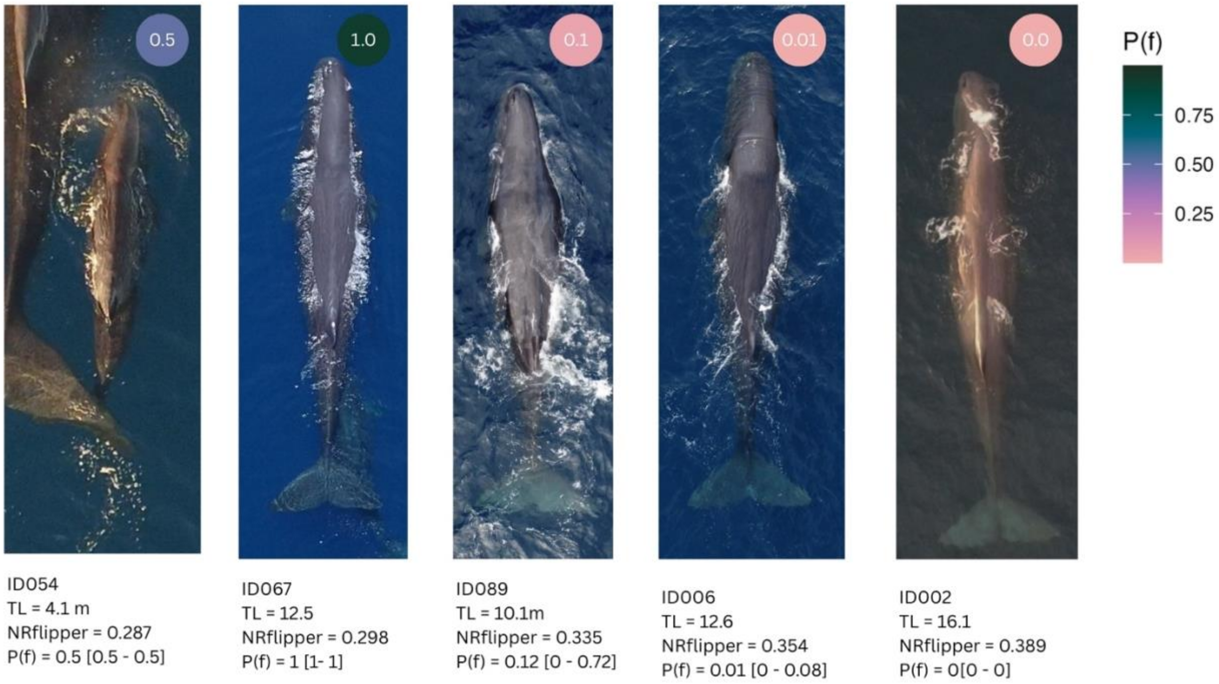


**Figure S4-1.** Sample still image close-ups of individual sperm whales analyzed. The circle at the top left of each photograph shows the bootstrapped mean posterior probability of that individual being female (*P(f)*). ID number, bootstrapped mean total length in meters (*TL*), mean $NR_{flipper}$ measurements, and the bootstrapped 95% CI range of *P(f)* are shown below each photograph. Individual ID054 is observed doing a peduncle dive onto another whale, and individual ID067 is shown receiving a peduncle dive

### References

Nishiwaki M, Ohsumi S, Maeda Y (1963) Change of form in the sperm whale accompanied with growth. Sci Rep Whales Res Inst Tokyo 17:1–17.

Pinheiro J, Bates D, DebRoy S, Sarkar D (2020) Nlme: linear and nonlinear mixed effects models. R Package Version 31-147.
